# Reexamining Waddington: Canalization and new mutations are not required for the evolution of genetic assimilation

**DOI:** 10.1101/2022.01.09.475581

**Authors:** Sarah Ruth Marzec, Katharine Pelletier, Amy Hui-Pin Chang, Ian Dworkin

## Abstract

Over 65 years ago, Waddington demonstrated ancestrally phenotypically plastic traits can evolve to become constitutive, a process he termed genetic assimilation. Genetic assimilation evolves rapidly, assumed to be in large part due to segregating genetic variation only expressed in rare/novel environments, but otherwise phenotypically cryptic. Despite previous work suggesting a substantial role of cryptic genetic variation contributing to the evolution of genetic assimilation, some have argued for a prominent role for new mutations of large effect concurrent with selection. Interestingly, Waddington was less concerned by the relative contribution of CGV or new variants, but aimed to test the role of canalization, an evolved form of robustness. While canalization has been extensively studied, its role in the evolution of genetic assimilation is disputed, in part because explicit tests of evolved robustness are lacking. To address these questions, we recreated Waddington’s selection experiments on an environmentally sensitive change in *Drosophila* wing morphology (crossvein development), using many independently evolved replicate lineages. Using these, we show that 1) a polygenic CGV, but not new variants of large effect are largely responsible for the evolved response demonstrated using both genomic and genetic approaches. 2) Using both environmental manipulations and mutagenesis of the evolved lineages that there is no evidence for evolved changes in canalization contributing to genetic assimilation. Finally, we demonstrate that 3) CGV has potentially pleiotropic and fitness consequences in natural populations and may not be entirely “cryptic”.

## Introduction

Canalization, a form of evolved organismal robustness, can facilitate the accumulation of hidden (“cryptic”) genetic variation (CGV)[1]. Waddington hypothesized that canalizing mechanisms and their evolution could facilitate rapid phenotypic evolution. As a test, Waddington experimentally showed that artificial selection for increased frequency of an ancestrally environmentally induced trait (loss of wing crossveins in *Drosophila*), resulted in the evolution of constitutive trait expression, a process he termed genetic assimilation[2]. Further work exploiting this system has shown that the evolution by genetic assimilation had a polygenic basis[3–5] apparently from segregating allelic variants[6] that interact with alleles influencing crossvein development[7]. More generally, CGV[8,9] and genetic assimilation[10–13] have been demonstrated in other contexts, and canalization, although not yet directly linked to the evolution of genetic assimilation, occurs in nature[14]. Although Waddington proposed genetic assimilation occurs through changes in canalization, alternative models have been suggested[6,15], and no explicit tests have occurred for canalization in this system. Similarly, the debate over whether genetic assimilation is due to segregating variation in the population or a result of *de novo* (new) mutations with large phenotypic effects [2,6] has reemerged in the context of high mutation rates of mobile genetic elements[16–18]. Additionally, recent work has demonstrated that in some instances, apparent CGV may actually not be phenotypic “cryptic”, but has previously unmeasured pleiotropic effects[19], and it is currently unknown whether this may be the case for the genetic variation contributing to the crossveinless system.

The increase in penetrance of the temperature induced loss of the crossvein under artificial selection is known to be due to a polygenic response, largely alleles of individually small effect[7,20]. However, several models have been proposed for the evolution of genetic assimilation. Waddington and others[6,21–23] discussed the relative contribution of standing genetic variation, both in terms of a polygenic response (many alleles of small or moderate effect interacting) and the contribution of rare alleles of large effect (i.e. “Mendelian” mutations). They also recognized the potential contribution of new mutations of large effect occurring concurrent with the selective response. In the polygenic response model, genetic assimilation occurs when the frequency of alleles across genes increase sufficiently that individuals (on average) have enough copies of alleles to result in a threshold effect (i.e. a liability-threshold model[6,15]). Alternatively, artificial selection can also select on rare segregating alleles of large phenotypic effects that cause crossveinlessness, facilitating genetic assimilation. Similarly, new mutations of large effect that arose concurrent with selection would produce similar results.

Early tests of these models favored a polygenic response based on standing genetic variation[3,6,7]. However crossveinless individuals are observed in natural populations at low (>1%) frequency, and spontaneous mutations in genes involved with crossvein development occur[4,24]. Furthermore, recent work has emphasized that heat stress[25,26], transposable element mobilization[27] and their interaction might increase mutation rates, and suggest new mutations are responsible for genetic assimilation[16,28]. Interestingly Bateman[6] examined the contribution of new mutations in 1959. Repeating the selection experiment using an isogenic strain (i.e. a strain with no variation), she observed no increase in frequency of the CVL phenotype nor genetic assimilation. Yet given current knowledge of transposable element mobilization rates under heat stress[29] this explanation has resurfaced.

We replicated Waddington’s experiment with modifications to augment aspects of the experimental design. After exposure to developmental heat stress each generation, individuals derived from a natural population were selected for loss of the posterior crossvein (hereafter “up-selection”), or its maintenance (“down-selection”). Unlike most previous studies (with limited replication) we generated six independent replicates of up-selection and three down-selection lineages, as well as controls for lab domestication to facilitate biological and statistical inferences. These lineages are used to answer questions as to the genetic architecture underlying the evolution of genetic assimilation for the crossveinless trait (i.e. to what extent, if any, CGV contributes to genetic assimilation), and whether canalization is necessary for the evolution of genetic assimilation.

## Results and Discussion

Consistent with previous findings, we observed a rapid response to selection for increased penetrance under developmental heat stress (Fig 1). We observed genetical assimilation in each up-selection lineage and from these propagated matching independent genetically assimilated lineages (Fig 1). Additionally, we generated three lab domestication lineages (with population sizes matching up- and down-selection) to account for laboratory adaptation and genetic drift.

**Figure 1:**
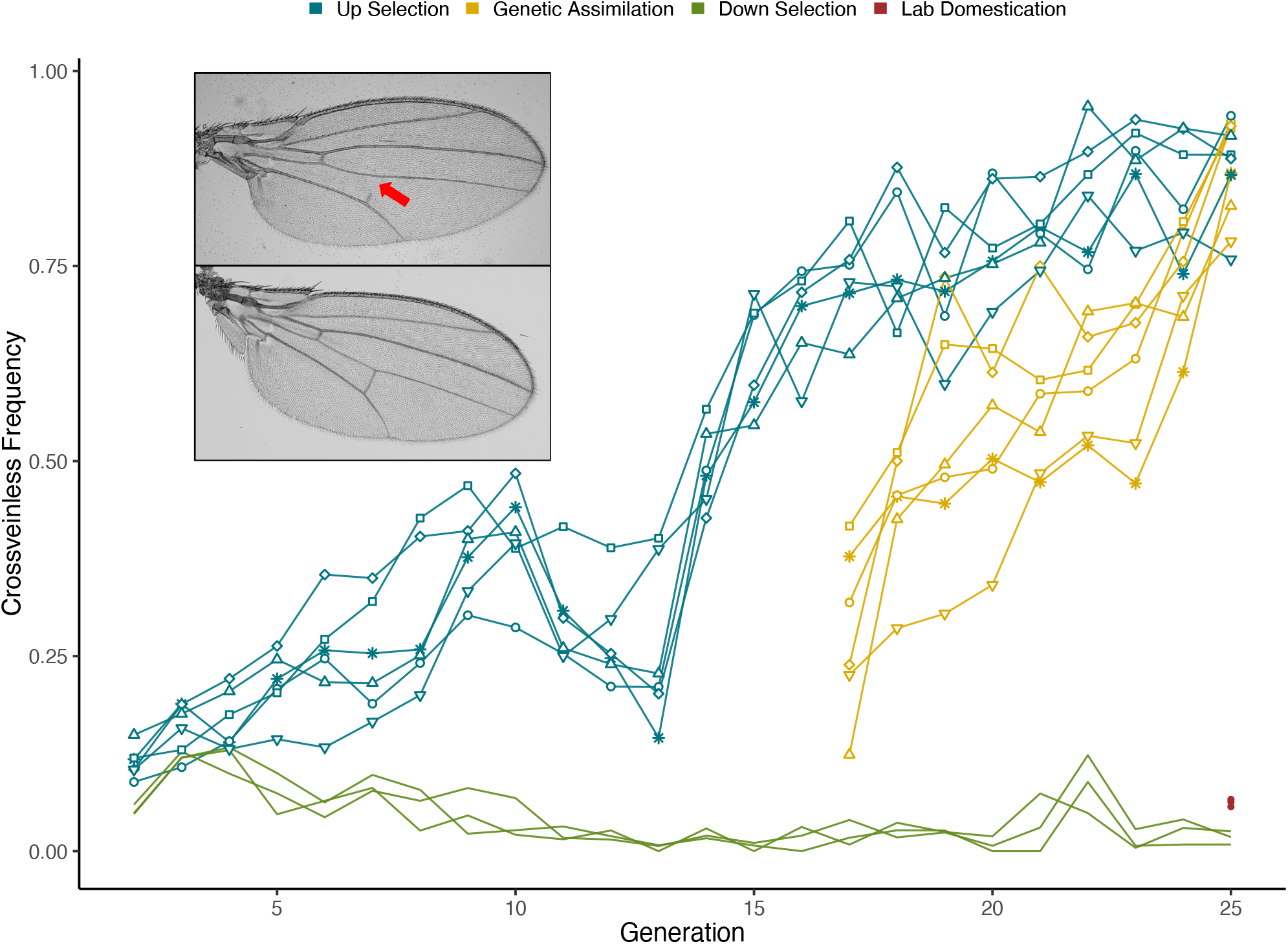
Alleles associated with CVL show strong response to selection. Crossveinless frequency is the proportion of flies for each lineage that showed the CVL phenocopy (defined as a fly with one or more breaks in one or both of the posterior crossveins, see picture) for that generation. Shapes on each up selection replicate lineage correspond to the matching genetically assimilated replicate lineage. Lab domestication lineages are shown only for generation 25 where they were heat-stressed to confirm maintenance of CVL alleles in these populations.

### Genetic assimilation of loss of crossveins is due to a polygenic response, not *de novo* mutations of large effect

To address the relative contribution of SGV vs new (or rare) mutations of large effect, we took multiple independent genetic and genomic approaches. Using crosses between all genetically assimilated lineages (and also some crosses between all up-lineages), we observed that F1 individuals from crosses between lineages showed only slightly reduced frequencies of loss of crossveins (S1 Fig). If the results were due to different new mutations in each independent lineage, then crosses would show highly reduced crossveinless penetrance, unless they had considerable dominance. However, if crossveinless is due to a polygenic response with partial parallel response of alleles, crosses would show intermediate penetrance similar to those of the “pure” lineages. Importantly, crosses of these lineages back to “control” (lab domestication) lineages showed low levels of crossveinlessness (S2 Fig), inconsistent with substantial dominance. Overall, the evidence is consistent with partial but incomplete parallelism among replicate lineages.

We next examined the potential role of genes known to influence crossvein development. Previous work suggests that distinct genomic regions contribute in different genetic assimilation experiments, often associated with genes influencing crossvein development[3,7]. However, when most of these studies of the genetic assimilation of crossveinlessness were performed, the number of known genes influencing this process was severely limited. Screening the Drosophila Flybase database we identified 81 genes with known loss-of-function (or RNAi) phenotypes influencing crossveins. We obtained co-isogenic control and deletion lines for 78 of these (spanning focal and nearby genes) and performed quantitative complementation analysis for all deletions by all six independent genetically assimilated lineages. This approach examines the relative dosage of allelic effects. If new variants of large effect contributed, we would expect few of these genes would fail to complement within each lineage (i.e. just the gene with the new mutation), likely with different genes interacting with each lineage. Under a polygenic response with partial parallelism we would expect many genes to interact with the replicate lineages and seeing partially shared response among the replicate lineages. We observed evidence that many, but not all, genes seem to be contributing (Figs 2 and 3, S3 Fig), varying among replicate lineages. Consistent with the results from the crosses amongst lineages, we observed evidence for polygenic response and incomplete parallelism amongst the assimilated lineages (i.e. Fig 4). Importantly, the deletions we examined had no effects on CVL frequency themselves under heterozygous conditions crossed to lab domesticated lineages, suggesting effects were not simply due to gene dosage. Our observation of incomplete parallelism (i.e. different genetically assimilated lineages in our experiment utilizing alleles in different combinations that ultimately result in assimilation of crossveinless phenotype) is consistent with SGV in that there is sharing of some groups of alleles. Parallel mutations occurring and rising to sufficient frequency in all lineages independently would likely require mutation rates at least 1000-10000 times greater than current estimates in *Drosophila*. Our results are also inconsistent with a single allele of large dominant effect as F1 heterozygotes of assimilated lineages have low penetrance (S2 Fig).

**Figure 2:**
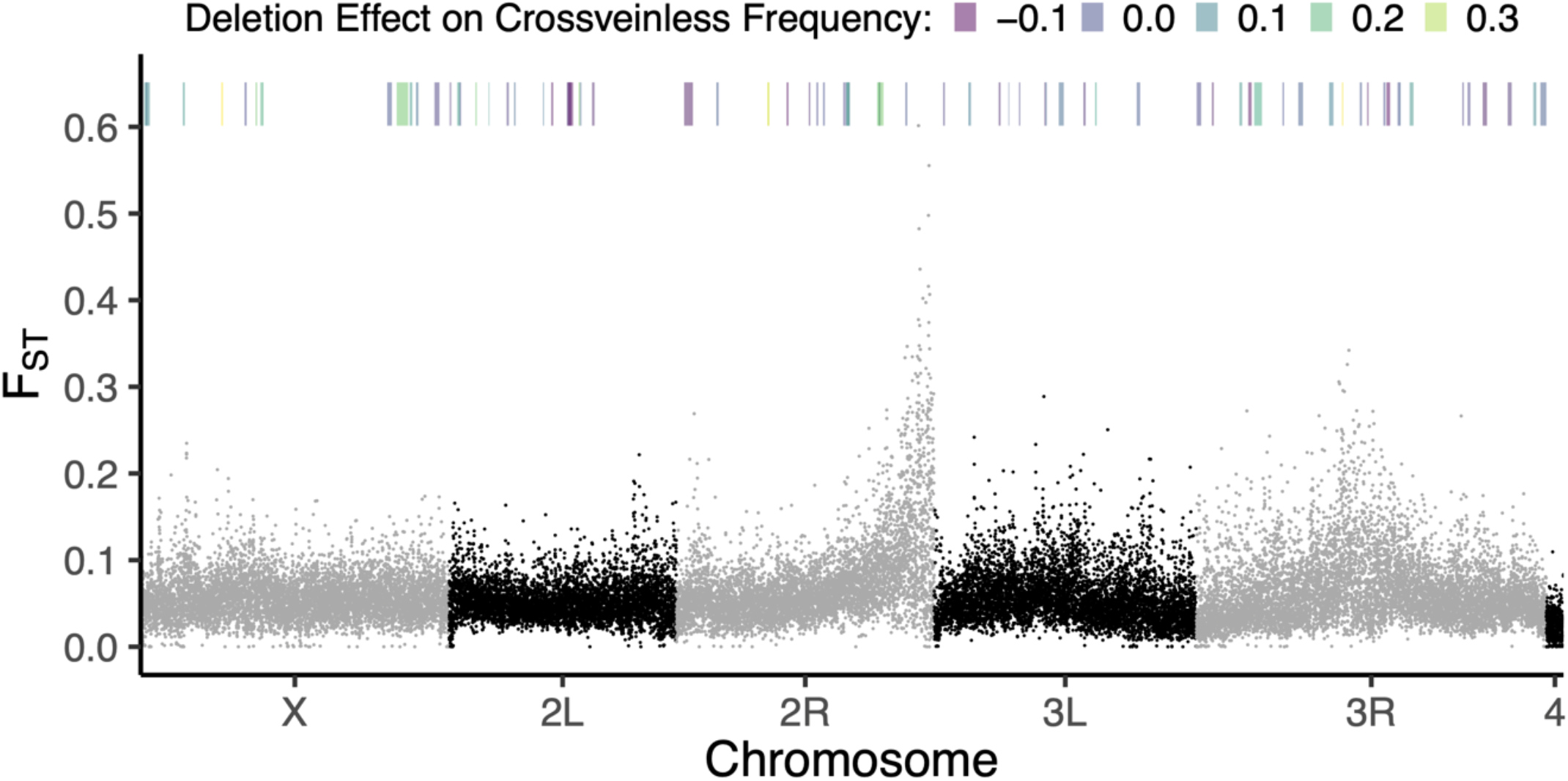
CVL phenotype is influenced by many alleles spanning the genome. F_ST_ was calculated using PoPoolation2 on 500 base-pair windows for comparing the genetic assimilation lineages with the ancestor population for two separate mapping software and taking the minimum F_ST_ value. The average effect of each deletion line influencing the CVL frequency in the genetic assimilation lineages (compared to control progenitor crosses) is depicted for the range of the genome that deletion spans.

**Figure 3:**
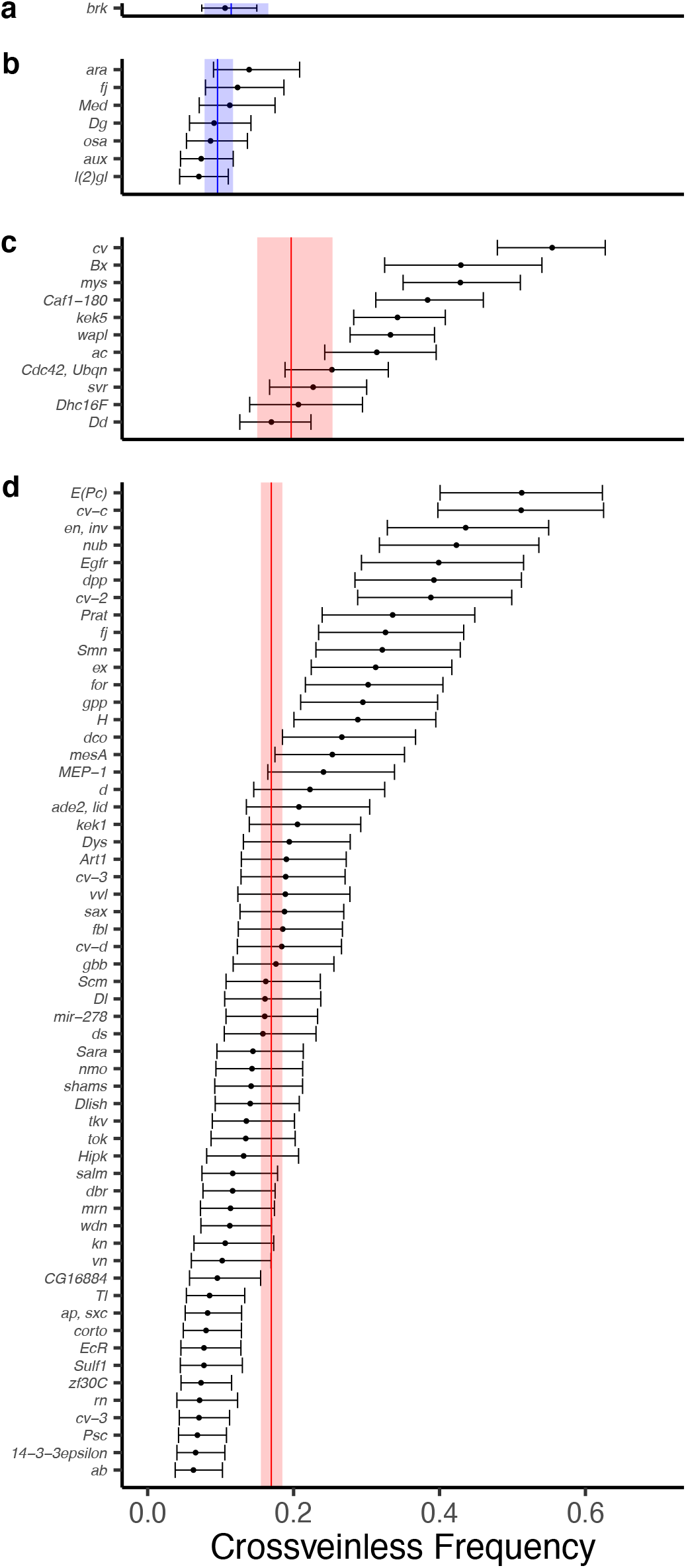
Deletion line crosses average for all genetic assimilation lineages. Genes of interest in each deletion region for (a) DrosDel X chromosomes, (b) DrosDel autosomes, (c) Exelixis X chromosomes, and (d) Exelixis autosomes (n=2 crosses for each deletion). Blue/red solid lines represent DrosDel/Exelixis progenitor means with shaded rectangles as 95% CIs. Error bars are 95% CIs on estimated effects from a generalized linear mixed model.

**Figure 4:**
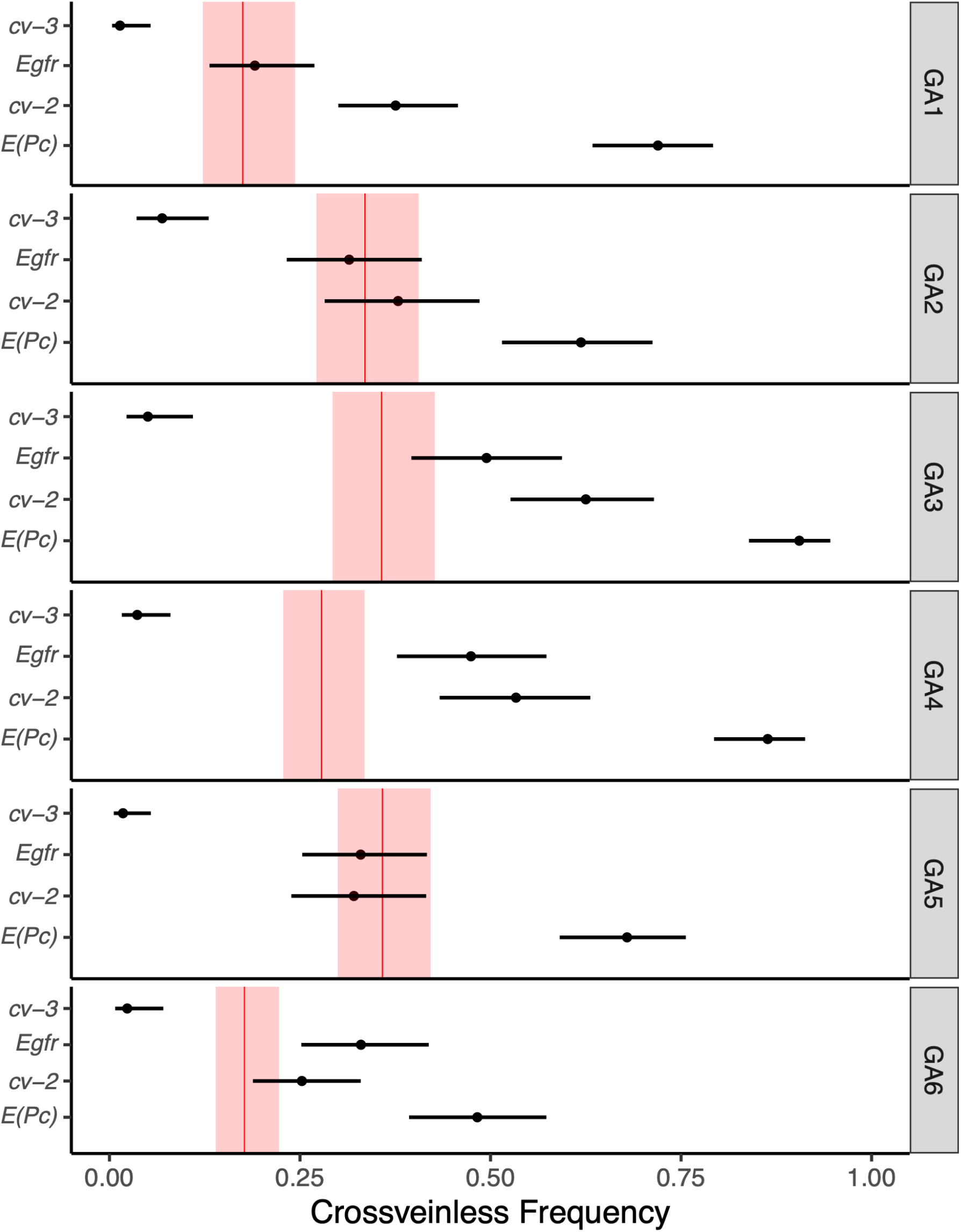
CVL phenotype in each lineage is uniquely influenced by alleles. Demonstrating a subset of deletion line crosses for each genetic assimilation lineage. Genes of interest in each deletion region for each genetic assimilation lineage (n=4 for each deletion). Red solid lines represent Exelixis progenitor means with shaded rectangles as 95% CIs. Error bars are 95% CIs on estimated effects from a generalized linear mixed model.

We used genome wide scans to examine the response in an unbiased manner (i.e. not dependent on candidate genes). If the response is polygenic due to a combination of segregating variants spread across many genes from in the ancestral population, we predict many small changes in patterns of genetic differentiation across the genome. Importantly this provides an explicit test of the model of new variants of large effect (or very rare segregating variants), even if their identity and position remain unknown. If genetic assimilation was due to new mutations of large effect followed by strong and rapid truncation selection (with the selected variant rising to near fixation), it would be associated with the signature of a hard sweep, substantially reducing nucleotide diversity in linked regions. We sequenced the ancestral and all selection lineages. We observed evidence of many regions in the genome contributing (Fig 2, S4 Fig), further support for a large polygenic response and not due to a small number of mutations of individually large effects. Genomic scans provided a strong test of the potential role of rare or *new* variants of large effect (S1 Table). While observing a small reduction in overall nucleotide diversity (relative to the ancestor) as expected due to drift, we do not observe large genomic regions of reduced variation indicative of a rapid hard sweep (Fig 5). We do observe smaller regions of reduced variation (S2 Table), and even those with small increases in a few genetically assimilated lineages. This is consistent with soft sweeps, polygenic responses and potentially some new variants of small phenotypic effects contributing and rising in frequency (but not yet approaching fixation). Overall, our results are consistent with a polygenic response, not new mutations (or rare segregating alleles) of large phenotypic effect, contributing to increased phenocopy penetrance and genetic assimilation.

**Figure 5:**
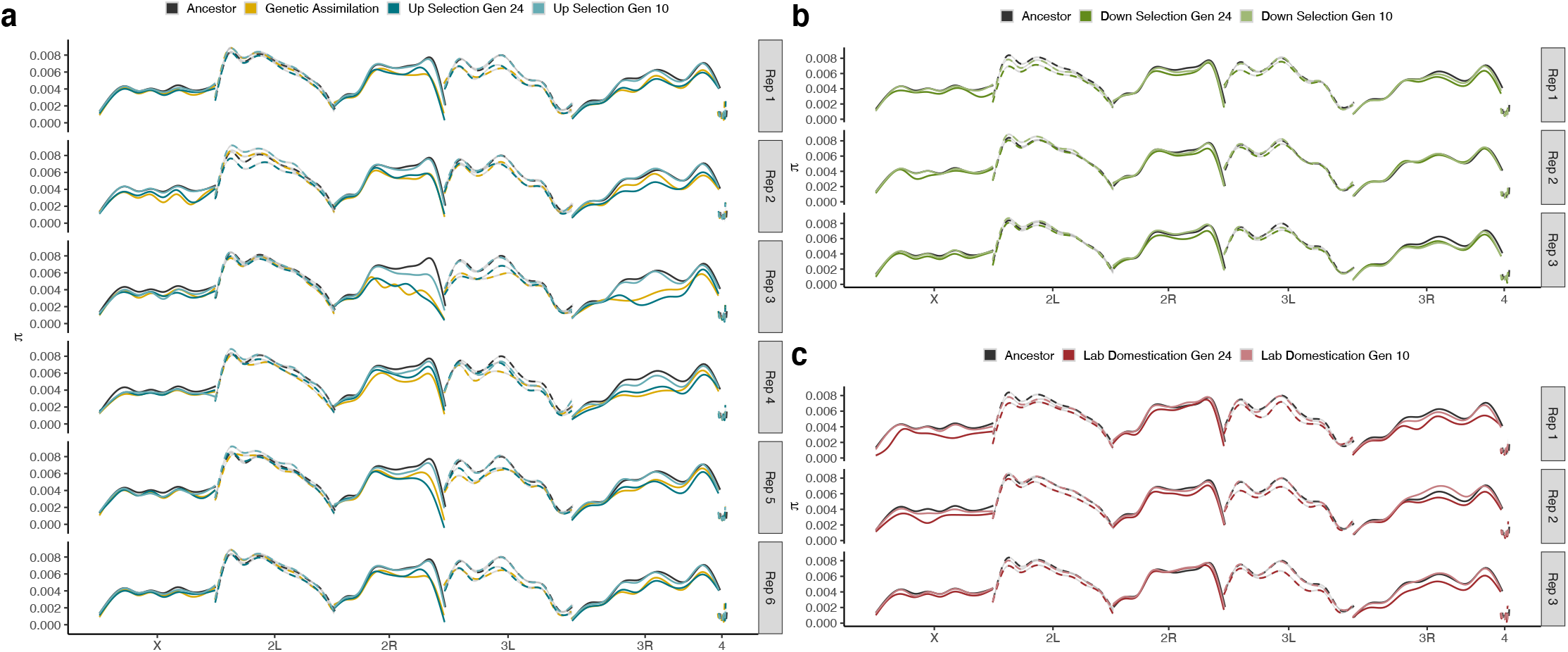
Nucleotide diversity (*π*) for up-selection lineages at two time points during artificial selection for (a) up-selection, (b) down-selection, and (c) lab domestication lineages. Nucleotide diversity was calculated on 500 base-pair windows for two separate mapping software and taking the minimum *π* value. Plots were generated with geom_smooth() using method ‘gam’ and formula ‘y~s(x,bs=“cs”)’ in R. Note that major regions of reduction correspond to centromeric and telomeric regions.

### Evolved Changes in canalization not needed for the evolution of a genetically assimilated trait

Perhaps the most contentious aspect of Waddington’s explanation for genetic assimilation is that its evolution required changes in the extent of canalization of traits to genetic and environmental variation[1,23]. Previous works suggests the canalization model was unnecessary, with genetic assimilation being explained as a threshold trait with underlying continuous genetic effects, i.e. a liability model[6,15] which is generally well supported, but which Waddington continued to reject as an explanation. Despite this, considerable work has examined conditions in which canalization can evolve, and several examples demonstrate variation in canalization[14,30]. Nevertheless, to our knowledge the role that canalization plays for the evolution of genetic assimilation specifically (and for CVL in particular) has not been appropriately tested.

There’s been many interpretations of the canalization model, but a common version is presented here (S5 Fig). If genetic assimilation is due to changes in canalization, we predict associated changes in sensitivity to perturbation. In our system, the constitutive classes (i.e. lab domesticated lineages with normal crossveins and genetically assimilated lineages with missing crossveins) should be the least sensitive compared with lineages with intermediate phenocopy frequencies during the evolutionary response (i.e. up-selection lineages). Waddington believed this should be true for both environmental and genetic effects, although these may in fact be independent evolutionarily[31]. As such we used independent experiments to examine whether genetic and/or environmental canalization evolved in a manner consistent with Waddington’s model.

With respect to genetic canalization, previous studies used release of CGV to infer canalization. Recent work suggests that examining the effects of new mutations is a superior method of investigation[32,33], which we employed here using mutagenesis. As the lineages were maintained as outbred populations, rare mutations of large effect segregating in the population need to be accounted for. As such we optimized the experimental crossing design, examined focal (wing) and control (eye) phenotypes as targets of mutagenesis, in addition to using non-mutagenized controls. Inconsistent with Waddington’s model, we did not observe an increase in mutational sensitivity in up-selection lineages compared with other evolved treatments. Additionally, lineages exposed to heat stress during artificial selection did not show higher levels of new mutations (in the absence of chemical mutagenesis), inconsistent with transposable element mobilization having a substantial impact on increasing genetic variation in these lineages (Fig 6, S6 Fig).

**Figure 6:**
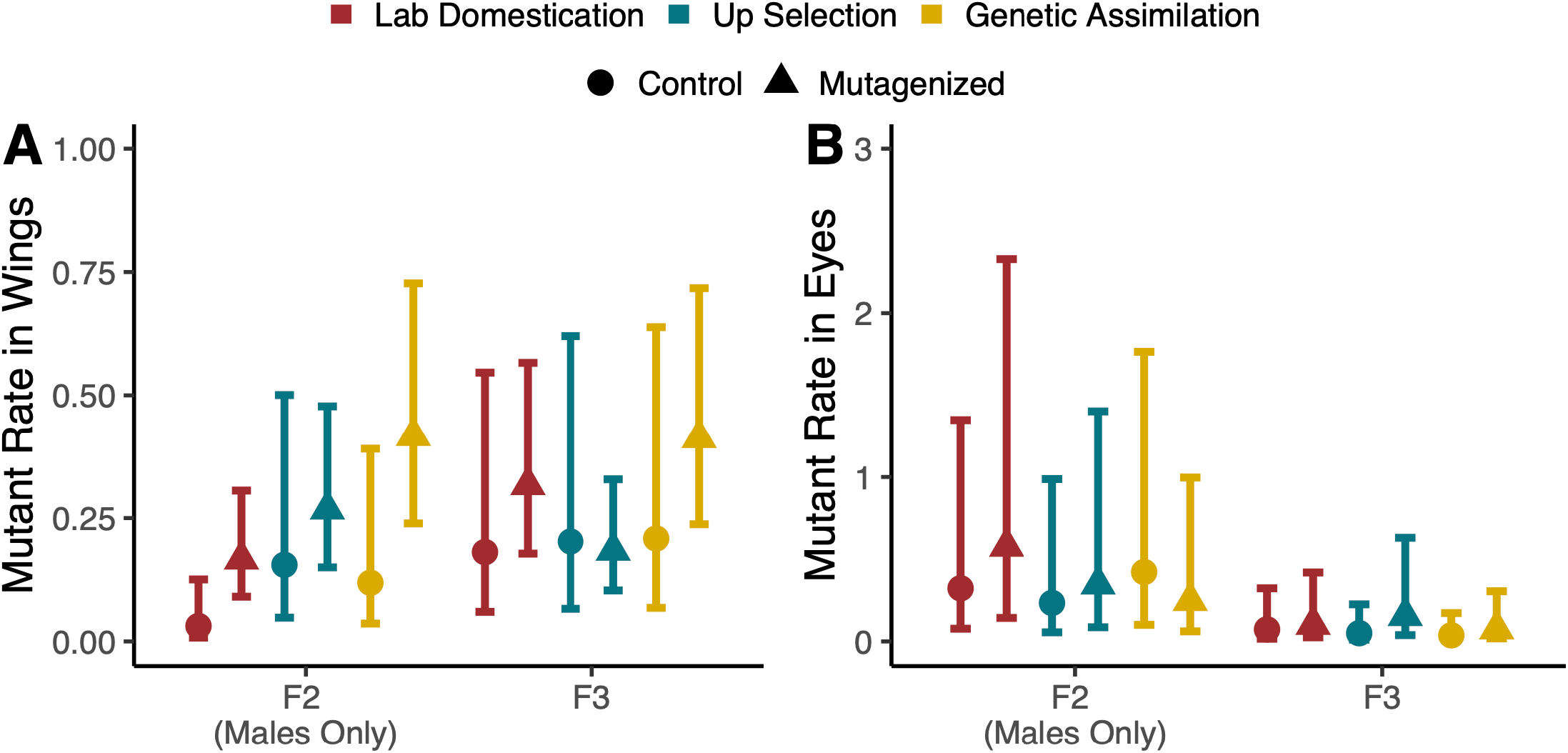
No differences among lineages for wing or eye defects induced by mutagenesis. Mutant rate is the estimated rate of observed phenotypes per individual for either (A) wings or (B) eyes. n_Mutagenized_ = 69-138 and n_Control_ = 24-30 replicate vials sorted for each selection lineage. Error bars are 95% confidence intervals on estimated effects from a generalized linear mixed model. In rare instances, single individuals showed multiple eye phenotypes which resulted in broad confidence intervals.

We examined whether these evolving lineages differed in robustness to environmental effects, in particular developmental temperature stress. We examined changes in mean and variance for wing size and shape as well as for qualitative wing defects (including crossveins). Mean size and shape changed with rearing temperature as expected, but with little evidence for differences in reaction norms among evolutionary treatments (Fig 7, S7 Fig). While our experimental design provides a weaker test for differences in micro-environmental variances, we also observed little difference among lineages.

**Figure 7:**
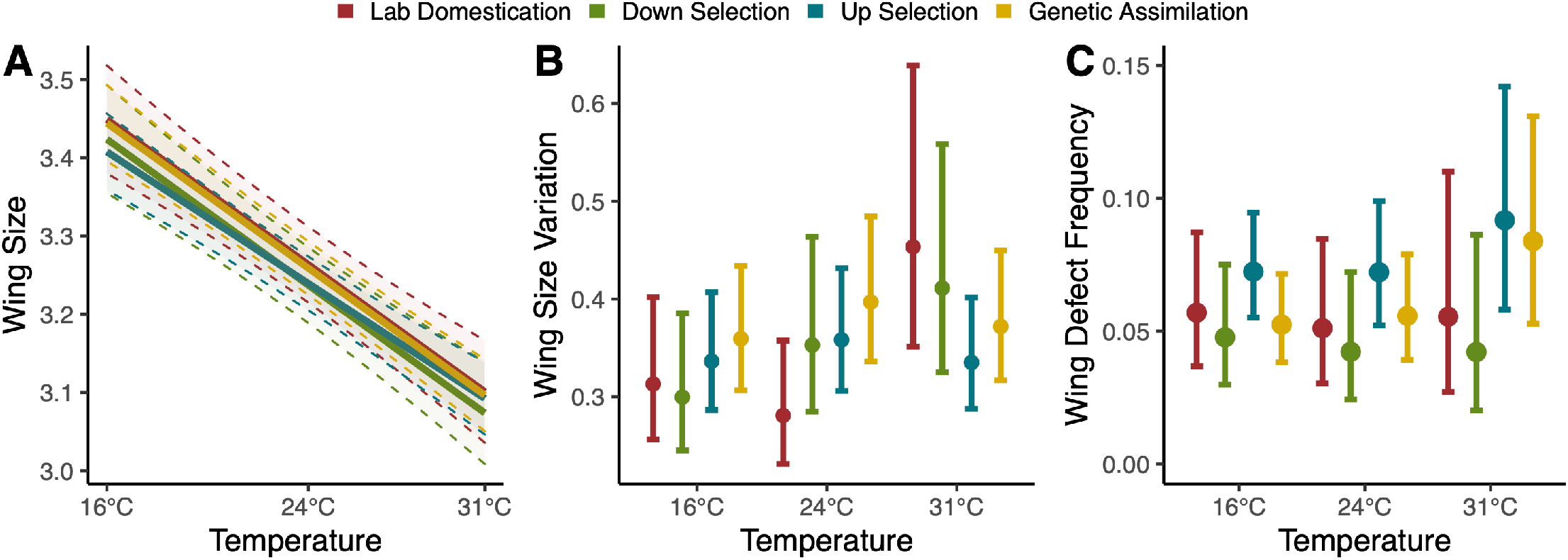
No differences in (A) macro-, (B) micro-environmental canalization, or (C) wing defect induced by rearing temperature. (A) Macro-environmental canalization measured as the slope of the temperature induced reaction norm for wing size (centroid size) among evolutionary lineages. Slopes are very similar with overlapping confidence bands. (B) Micro-environmental canalization measured as wing size variation (using Levene’s statistic) within selection lineages for different rearing temperatures. Error bars are 95% confidence intervals on estimated effects from a generalized linear mixed model. ANOVA shows treatments do not differ from each other (p>0.7). (C) Wing defects include anterior crossvein, wing margin, and longitudinal vein defects. All defects were combined for modeling due to rarity of defects. Error bars are 95% confidence intervals on estimated effects from a generalized linear mixed model; none of the treatments show significant differences from each other at either density (p>0.1).

These results demonstrate that neither genetic nor environmental canalization have evolved among our evolutionary treatments, despite rapid evolution for genetic assimilation of crossveinless wings. Thus, in short time scales (less than 25 generations), changes in canalization are not necessary for the evolution of genetic assimilation nor is there a substantial role of new mutations of large effect in the genetic assimilation response. While canalization clearly can and has evolved[14], its role in the process of genetic assimilation may not be as relevant. Similarly, the role of TEs in generating variation, including adaptive effects is clear[34], but it may have a minor contribution in the evolution of GA relative to a polygenic response based on standing genetic variation.

### Allelic variation contributing to selective response may not always be “cryptic”

If the response to selection is a result of standing genetic variation in natural populations, are the allelic effects truly “cryptic” in environments without temperature stress? If so, the expectation would be that their frequencies in natural populations are maintained under mutation-drift balance. Alternatively, such allelic variants have unmeasured pleiotropic effects[19], and are maintained in the population in part due to selection. In this scenario we would predict a correlation between response to selection for loss of crossveins and other traits or fitness. Utilizing these lineages, we sought to determine the aggregate fitness consequences of alleles contributing to the response. A previous study suggested that the alleles contributing to the CVL phenocopy response were potentially deleterious[6] by examining how phenocopy penetrance varied under relaxed selection. Using a similar form of relaxed selection with additional treatments for with and without heat stress, we observed a decrease in CVL frequency relative to the starting generation (S8 Fig), consistent with a deleterious effect of these alleles in aggregate in up-selection lineages.

If alleles contributing to the crossveinless phenocopy response are maintained by selection in natural populations, we may see contrasting effects on fitness components. We examined three fitness components of *Drosophila*: viability, fecundity, and male mating success. The up-selection lineages showed reduced viability compared with the down-selection lineages at low density, with no significant differences among selection lineages at high densities (Fig 8a). To confirm those effects we performed a subsequent experiment examining competitive viability and observed consistent effects (Fig 8b). We also examined both fecundity and male mating success in parallel experiments. While there was considerable variation, we did not see substantial treatment effects (S9 Fig). These results suggest that -- in aggregate -- alleles that contribute to this response may in fact be deleterious independent of heat stress, at least in these genetic backgrounds and conditions.

**Figure 8:**
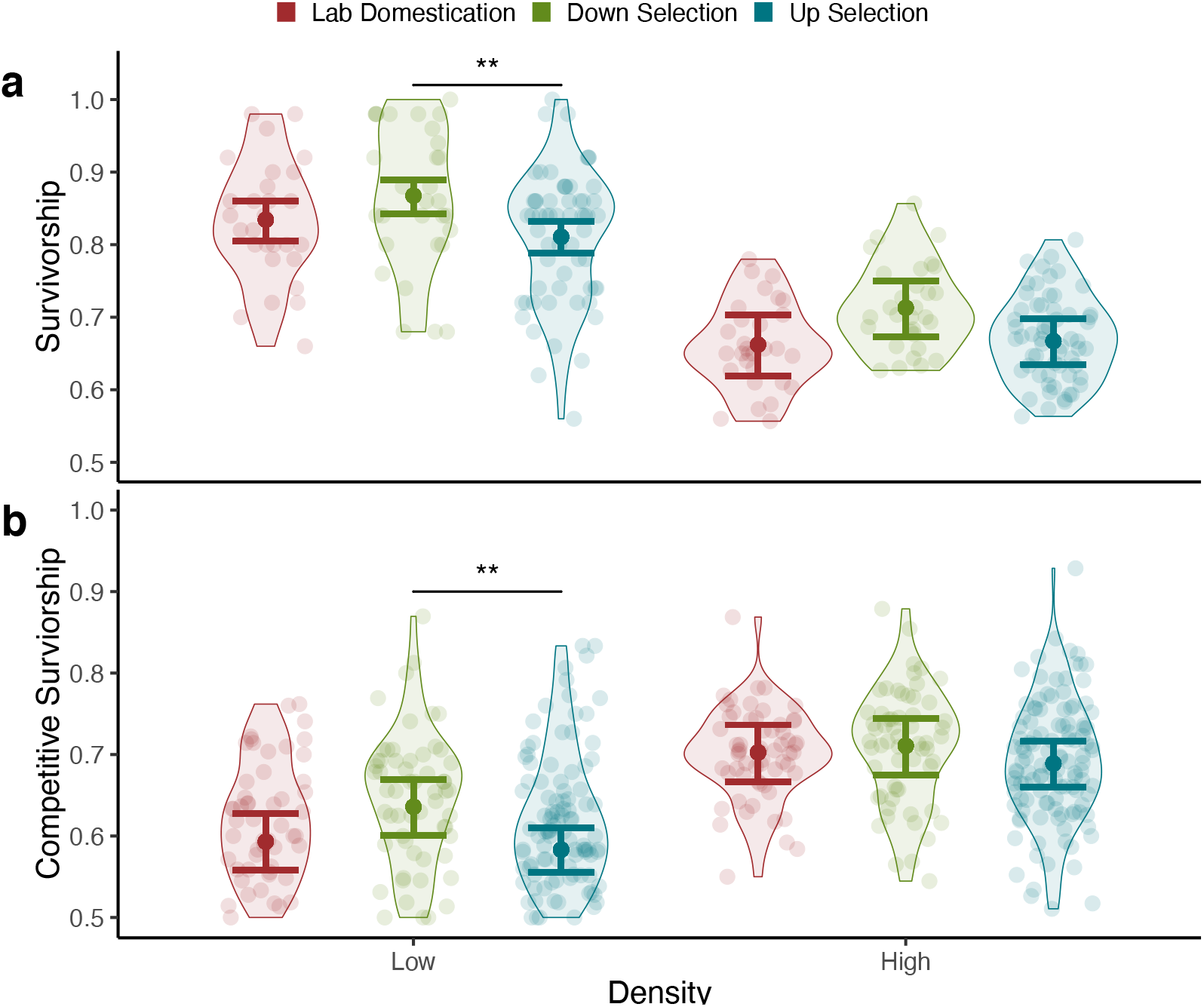
Increased frequency of CVL alleles in a population is associated with reduced viability. Percent survivorship of the three selection regimes when (a) alone or when (b) half the eggs are a common competitor. High density had 300 eggs and low density had 50 eggs per vial. For each density: n_LD_=30, n_DOWN_=30, n_UP_=60 vials (10 vials per replicate lineage). Transparent points show variation within lineages and opaque points are fitted values for each lineage. Error bars are 95% confidence intervals on estimated effects from a generalized linear mixed model (**P < 0.01).

We also examined whether these alleles have correlated or pleiotropic effects on specific traits. The number of measurable traits is impossibly large, but using the 83 genes in Flybase with identified roles in crossvein development, gene ontology enrichment suggests that many have additional roles during organismal development including cell proliferation, body size and other aspects of wing development. Indeed, wing size, shape, and development are known to be influenced by variants in genes that contribute to crossvein development[35–37]. While there was considerable variation in body size, wing size and wing shape we observed no evidence for consistent treatment effects on mean trait values (S10 Fig).

While we were unable to demonstrate what specific traits the fitness effects are linked to, our results suggest that the allelic variants influencing crossveinless phenocopy response influence fitness in aggregate, and are not necessarily phenotypically “cryptic”. As this trait is highly polygenic, it could be that we are underestimating phenotypic effects as the pleiotropic contributions of alleles influencing CVL could vary in direction. Further, it may be that the fitness effects of the alleles influencing viability are maintained due to other forms of selection (i.e. density or frequency dependence). It should be noted that the allelic combinations brought about by selection may be rare in nature, and thus the phenotypic effects could be largely cryptic in such circumstances.

### Insights

In readdressing Waddington’s classic experiment, we have found that evolution of genetic assimilation is due to a polygenic response largely from segregating alleles and we find no evidence for the contribution of changes in canalization during this process. Furthermore, the CGV contributing to this response may not be entirely phenotypically “cryptic” and may contribute to variation in fitness in natural populations under some circumstances.

While canalization is not necessary for genetic assimilation, it does occur in natural systems[14,38,39]. Similarly, while new mutations of large effect due to increased mobilization of transposable elements is not relevant here, their role in adaptive response in nature has been demonstrated[34]. While we demonstrate that the variation influencing the CVL phenotype may not be phenotypically “cryptic” in all circumstances, we note that the fitness effects we observe may be due to allelic combinations rarely occurring in nature. Since we observed aggregate effects of a polygenic CGV response, we cannot disentangle the individual phenotypic effects and it is likely that they may cancel out if they differ in sign. Future work plans to address the polygenic nature of this response and identify specific causal variants and possible genetic interactions amongst them.

As with other seemingly puzzling phenomena, extraordinary claims about mechanisms contributing to genetic assimilation have been proposed, often with tenuous evidence. In particular, the contribution of epigenetics, canalization, and high rates of transposable elements mobilization have been recently emphasized as alternatives to the role of standing genetic variation contributing to genetic assimilation. However, following the Sagan Standard[40,41], “extraordinary claims require extraordinary evidence”. In the case of genetic assimilation of CVL, both current and past evidence is consistent with a “typical” polygenic response on an underlying threshold trait primarily due to standing genetic variation. While for a lab based artificial selection experiment addressing questions of genetic assimilation and canalization this may seem academic. However, given the increasing need to evaluate how populations and species may survive rapid anthropogenic changes, any potential roles of CGV and genetic assimilation in contributing to evolutionary rescue become increasingly important. These mechanisms may not only be less parsimonious but also detrimental to our understanding in how to care for populations responding to climate change and undergoing rapid evolution.

## Materials and Methods

### Establishment and Maintenance of *Drosophila* Selection Lineages

Wild *Drosophila melanogaster* were collected at Country Mill Orchard (Michigan, USA 42°38’8.5”N 84°47’32.3”W) in Fall of 2015. Field collected females were individually placed into vials so that male offspring could be used to distinguish *D. melanogaster* from *D. simulans*. Two male and two female progeny from each field collected *D. melanogaster* female were used to generate the lab population. Progeny from over 1000 field collected D. melanogaster were used to establish this population. Importantly we purposefully used a freshly collected population (as opposed to a lab domesticated population) because of the potential critical role of standing genetic variation in natural populations. After establishing the base population individuals were stored in 70% ethanol (−20°C) for future genotyping. All flies were maintained using standard cornmeal media (recipe https://github.com/DworkinLab/Protocols/blob/master/Recipes.md).

The population was maintained for 3 generations until being split into replicates for each selection regime: a control group subject only to lab conditions, representing a control for lab adaptation (***lab domestication***), as well as selection regimes exposed to high temperature stress (***up-selection*** and ***down-selection***). There were a total of 12 replicate selection lineages: 6 up-selection replicates, 3 down-selection replicates, and 3 lab domestication replicates. Each replicate lineage was initiated with 50-pairs of flies and allowed to mate for one day in cages (bugdorm 17.5cm^3^) maintained at 21°C. Flies laid eggs over the next five days, with one bottle switched out once a day to maintain low to moderate larval density, maintaining genetic diversity. Cultures in bottles were maintained at 24.5°C. Pupae for the up-selection and down-selection lineages were heat-stressed (described below) and then returned to 24.5°C. Eclosed flies were sorted daily for sex and kept at 18.5°C to allow for minimal development gaps between the flies eclosing early/later in the week. The selection regime described was completed on a three week cycle for each generation, where week 1 was egg-laying, week 2 was for pupae collection and heat-stress exposures for the up-selection and down-selection lineages, and week 3 was for collecting and sorting adults. This was repeated for each generation of selection.

### Establishment of artificially selected lineages

#### Staging of pupae and high temperature exposure

The critical window for the temperature mediated crossveinless phenocopy occurs during early-mid pupal development[5,21] [stage P5 from [42]]. We used standard procedures to procure staged cohorts of *Drosophila*. Pupae develop an air bubble causing them to float to the water surface at 8 hours past pupation if developing at 24.5°C. Pupae from media bottles were collected and age of pupae were estimated using a series of two floatings in water. Pupae aged at 8±2 hours past pupation were retained to be used for temperature experiments. Based on previously published work[20] and pilot experiments to examine both the penetrance of the crossveinless phenocopy as well as viability, pupae were exposed to 37.5°C for four hours at 24±2 hours past pupation. This approach differs from Waddington’s in that we used a longer temperature stress but a lower temperature. We made this changes as Waddington’s original procedure had a substantial impact on viability which would cause a strong selective response in its own right as well as contribute to increased genetic drift, both of which could confound analysis and interpretation of experiments. Our collection of pupae from several replicate lineages necessitated a ±2 hour window on pupae age, and a longer duration of heat stress increased the overlap with the critical development stage for the majority of the pupae. The length and magnitude of the temperature shock we used are similar to numerous other studies which examined aspects of the crossveinless phenocopy[3]. Heat-stress exposure was performed by placing pupae on moistened paper-towel in plastic vials with plugs. The vials were submerged to just below the top of the vial in waterbaths within a plastic rack.

Virgin adults were sorted for crossveinless phenotype defined as having at least one break in the posterior crossvein of either wing. Up-selected flies were selected for having crossveinless wings while down-selected lineages were selected for having wild type crossveins. In the first few generations, some replicate lineages did not have 50 pairs of individuals with loss of crossvein (up-selection only), and were supplemented with individuals from the same replicate. This was done to make sure that population size was not reduced. Individuals from the lab domesticated lineages were randomly selected at the same time. 50-pairs were chosen from the selected flies for each replicate lineage to initiate the next generation. Parental flies from each generation were stored in 70% ethanol at −20°C.

#### Generation of Genetically Assimilated Lineages

Genetically assimilated flies (crossveinless flies that developed ***without*** high temperature stress) were first examined and observed in generation 15. Starting in generation 17, a subset of pupae from each up-selected replicate lineage were allowed to develop without high-temperature stress. Crossveinless flies were selected to start matching assimilated lineages. These assimilated lineages were maintained separately from their corresponding up-selected lineages (and from other assimilated lineages) but were supplemented with additional assimilated flies from their corresponding up selection lineages for the beginning generations since we could not find 50 male-female pairs per genetically assimilated lineage at the start (S3 Table) to maintain equivalent census population sizes. By generation 3 of assimilation (generation 20 of selection lineages), lineages no longer needed to be supplemented. Assimilated lineages were maintained with selection, but without heat stress.

### Estimating fitness effects

#### Relaxed Selection on phenocopy penetrance

At generation 18, each of the six replicates of up-selected lineages were split into three sets (treatments). One set acted as a control and continued the normal high-temperature stress and selection protocol. The second set was exposed to high-temperature stress, but no selection was performed. The third set had neither exposure to high-temperature stress nor selection. Each set of lineages were otherwise maintained normally for five generations. Progeny of the fifth generation (parallel to generation 23 of main experiment) were exposed to high-temperature stress and CVL frequencies determined by counting 100 flies /replicate/sex and modeled using a logistic mixed model with treatment as fixed effect, and random effects of lineage accounting for repeated measures using glmmTMB (v0.2.3). All analyses were performed in R (v3.5.0).

#### Fitness component assays

Fitness assays were done with subsets of individuals from lineages unexposed to temperature stress for two generations prior to experiments to avoid confounding maternal effects of heat stress. Assays were done with all replicate selection lineages except assimilated.

##### Viability

Individuals were split from selection lineages at generation 28. Eggs from each selection lineage replicate were placed in vials of low or high (50 vs. 300 eggs) density. We used both densities to mimic natural as well as lab evolved conditions. 10 replicate vials per density/replicate lineage within each evolutionary treatment were used. Viability was measured as proportion of surviving adults. A logistic mixed model was fit with treatment, density and their interaction as fixed effects. Independent random effects were fit for collection date, individual (egg picker), and replicate lineage nested within treatment, including “random slopes” for density. This was done in lme4 (v1.1-19) using glmer(), and Anova() in car (v3.0-2).

##### Competitive Ability

Eggs from each replicate selection lineage were placed in vials at either low or high density (50/300 eggs) with half the eggs from a marked competitor; the recessive *scute*^*1*^ allele introgressed into the ancestral background population. There were 10 replicate vials/density/replicate treatment, split from selection lineages at generation 38. This was repeated with a second block in generation 45. Competitive viability was analyzed in a similar manner to viability, with the addition of uncorrelated random effects for experimental blocks and effects of “person” transferring common competitors to vials.

##### Fecundity

Females (n=21-24) from each replicate selection lineage (lab-adapted, up-selected, and down-selected) and larval density were collected from the viability assay. Each female was placed in a vial and mated for 24 hours with a male from the same treatment/density. The pair of flies was then flipped into fresh vials each day for 5 days. At the end, all five vials were collected and total eggs counted for each female. Switching the female into a new vial each day facilitated egg counting and mirrored artificial selection protocols, where bottles were transferred daily. Female thorax length was imaged (Leica MZ12.5 microscope, 6.3x magnification) and measured (ImageJ v1.50f) to include as a covariate as size influences female fecundity. A linear mixed model was fit with treatment, density, their interaction and thorax size as fixed effects, and replicate lineage nested within treatment as a random effect using lmer().

##### Competitive Mating Ability

Individuals were split from selection lineages at generation 40. The marked competitor population with *scute*^*1*^ mutation was used because it was easily distinguishable and had no other previously known fitness effects. A male from each treatment lineage and a *scute* male were placed in a vial with a single *scute*^*1*^ female (n=37-50). Replicate vials were set up in a balanced block design with equal numbers of lineage replicates. Females were allowed to lay eggs for 3 days then all three flies were switched to a new vial to keep density low. Progeny were counted and sorted for *scute*^*1*^, totaled over 6 days, to estimate proportion sired by each male. A logistic mixed model was fit with treatment as a fixed effect. Independent random effects were fit for replicate lineage nested within treatment, vial replicate nested within replicate lineage and treatment, and block using glmer().

#### Pleiotropic (correlated) effects of CVL alleles on body size, wing size and shape

Using the flies from the fecundity assay described above, thorax length was used to examine correlated effects of CVL selection on size. A linear mixed model was fit with treatment, density and their interaction as fixed effects, with random effects for replicate lineages within treatment for both intercept and density. Animals were stored in 70% EtOH until right wings from females (low density only) were dissected, mounted in 70% glycerol in PBS and imaged (Olympus DP80 camera mounted on an Olympus BX43 microscope, using a 4X objective total 40X magnification). Images were captured with cellSens Standard (V1.14) at 4080 × 3072 pixels (0.0005375 mm/px). Landmarks were obtained using Wings (v. 3.72) and CPR (v1.11)[43] to extract Procrustes superimposed configurations with 12 landmarks (excluding posterior crossveins) and 33 semi-landmarks plus centroid size. 480 individuals (average 20/vial/replicate), were included. To examine correlated effects on wing size we used the same model described above for thorax length, without effects of density. The median form of Levene’s statistic was used for variability in wing size (examined on both linear and log scale). A similar model to that described above was used, but fit using an inverse link function and assuming Gamma distributed error with glmmTMB().

Analysis of wing shape was done in geomorph (v. 3.1.3)[44], with fixed effects of centroid size, treatment, the interaction between treatment and replicate lineage and the interaction between treatment, replicate lineage and vial replicate on shape residuals. For hypothesis testing, a model without the treatment effect term was used as the null model. The effect of treatment for both mean shape (using distance between vectors) and variance (using disparity) was tested with these models.

In a second run of this experiment, we addressed whether there were differences between the up-selection and their corresponding assimilated lineages for patterns of variability. For this experiment, we used flies saved from the “Relaxing Selection on Flies” were used for non-heat-stressed up-selected lineages along with lab-adapted lineages from the matching generation (F23) and genetically assimilated flies of the corresponding generation (F6). Wings from female flies were dissected and shape data was collected as described above. A total of 291 individuals were included, averaging 19.5 individuals for each replicate lineage. The effect of treatment on wing size was tested using a mixed model with a fixed effect of treatment and a random effect of replicate lineage. To test the hypothesis that treatment has an effect on wing size variance, a generalized linear model was used with the same predictors as the size model above. For shape, a model with terms for centroid size, treatment and replicate lineages was fit using the geomorph package. For hypothesis testing of the effect of treatment on mean shape change and variance, the null model removed the treatment term.

### Isolation of DNA

Isolation of DNA was done with a modified Qiagen kit protocol for “Purification of total DNA from insects using the DNeasy Blood & Tissue Kit - using a mortar and pestle”. 100 individuals from each replicate lineage were used in the DNA extraction (50 males and 50 females) for pooled sequencing. Flies were prepared in groups of 25 individuals (by sex). After DNA quantification, the four groups making up a selection lineage replicate were combined to have equal DNA contributions. The ancestral population was sampled and sequenced with a total of 400 individual flies to capture genetic diversity and rare variants in the founding population, but with the same group (25) and pool (100) sizes.

### Genomic Analysis

To examine the genomic consequences of selection and assimilation, we sequenced all up-selection, down-selection, and lab domestication replicate lineages at generation 10 and generation 23. Genetic assimilation lineages were sampled at generation 8, corresponding to generation 23 of other selection lineages. This was done at the Michigan State University RTSF Genomics Core using the Illumina Truseq Nano DNA library preparation kit and samples were run over two days on 4 lanes (4×2 lanes total) Illumina HiSeq flow cell. We generated 125bp paired reads with an average insert size of 700bp. We obtained ~140X genome coverage for each selection lineage replicate (at each time point) and 600X coverage for the ancestor population. Reads were mapped to the *D. melanogaster* reference genome (r5.57) using bwa (v0.7.8) and novoalign (v3.07.00). We used two mappers which had both been evaluated for pool-seq data and used the intersection to reduce false-positive polymorphisms [recommended in [45]]. PCR duplicates were removed with Picard (v1.131) and GATK (v3.4-46) for indel realignment. Nucleotide diversity (*π*) and F_ST_ were calculated with PoPoolation (v1.2.2) [46] and PoPoolation2 (v1.201) [47], respectively (--pool-size set to individuals per replicate, --max-coverage set to approximately double the average genome coverage per replicate).

### Crosses among lineages

Reciprocal crosses were performed among selection lineages, and CVL phenotype frequency determined in F1 progeny and compared to frequencies of corresponding “pure” replicate lineages. Heat stress was applied as described above. Up-selected lineages were crossed in a balanced incomplete round-robin design due to constraints of having to heat-stress pupae. We performed each cross at two generational time-points (corresponding to generations 22 and 23) and at least 100 flies were scored for each sex/cross.

For genetically assimilated lineages all possible crossing of lineages were done, but otherwise the details of the crosses (but without heat shock) are as described above. We performed each cross at two generation time-points (corresponding generations for assimilated lineages are 12 and 13) and at least 100 flies were scored for each sex per cross. To quantify differences between pure lineages and “hybrids” between them for both the up-selection and assimilation crosses, a logistic mixed model was fit using CVL counts, with pure/hybrid and block as fixed effects, and independent random effects for lineage from both male and female parents.

### Quantitative Complementation Test - with co-isogenic deletion lines

Using FlyBase (FB2017_06), we identified the ~81 known genes influencing crossvein development via loss-of-function (mutant or RNAi) perturbation. Using this set, we identified deletion lines within each of the Exelixis and DrosDel Deficiency collections. While each of these collections had independent progenitor strains, all deletions within collection are otherwise co-isogenic to their respective progenitor. We identified 76 deletion lines, spanning 78 genes of interest. Every genetically assimilated replicate lineage was crossed with each deletion line (n = 2 independent crosses) and its respective progenitor control lineages (n = 2 crosses per block). Given the large number of experimental crosses required (>900) multiple experimental blocks were required, with independent control crosses within each block. Males of autosomal deletion lines were crossed with females of genetically assimilated lineages and all progeny containing the deleted chromosome region were scored for CVL frequency. Females of X-chromosome deletion lines were crossed with males of genetically assimilated lineages and female progeny were scored for CVL frequency. Twelve deletion lines were further tested with a higher number of replicate crosses per lineage (n=4). Measures of CVL frequency were done as stated above.

To confirm that the effects observed were not due to haploinsufficiency of the deletion, Males from 10 autosomal deletion lines (and females from 2 X-chromosome deletion lines) were crossed to 3 lab-adapted lineages (LD1: n = 2 vials, LD2: n=1, LD3: n=1). Among F1 progeny, no individuals with the crossveinless (CVL) phenotypes were observed.

To serve as a background matched control to examine dominance of the CVL phenotype we crossed each genetically assimilated lineage with the 3 lab-adapted lineages reciprocally (n = 4 independent crosses each) and CVL frequency recorded among F1. For each deletion we fit a logistic regression (counts of CVL and wild-type flies) with genotype (deletion/wild type), assimilated lineage and their interaction. For several sets of crosses we observed complete separation during data modeling. As such we adjusted all counts by adding one to CVL and one to wild-type to enable model convergence.

### Mutagenesis (Genetic Canalization)

See S11 Figure for graphical representation of methods. Males (G0) from three replicate lineages of the up-selection, lab domesticated, and genetic assimilation selection treatments were starved for 12 hours (with wet cotton ball for hydration), exposed to 25mM EMS (in 1% sucrose solution) for 8 hours and allowed to recover for 24 hours. After, they were mated to virgins of the same lineage. Along with the mutagenized males, control (non-mutagenized) males were fed with a 1% sucrose solution and then mated with virgin females of the same lineage (Mutagenized males n=23-46; Control n=8-10 from each lineage). F1 progeny were split into 3 single pair crosses from each G0 sire. F2 males (for X-linked mutations), and F3 males and females were scored for eye {color, shape, size, roughness} and wing {shape, size, scalloping, curling, crumpling, pigmentation and venation including anterior crossveins} phenotypes. The total number of individuals scored was 60,638.

A second mutagenesis experiment was performed to additionally examine the effects on down-selection treatment and to examine longer EMS exposure (16hrs) using males from three replicate lineages of each selection treatment (up-selection, down-selection, lab domesticated, and genetic assimilation). G0 males were mated to virgins as above (Mutagenized n=9-10; Control n=5 for each). F1 progeny for each were split into 2 single pair crosses. Phenotypic scoring as described above. The total number of individuals scored was 40,827.

A logistic mixed model was fit with mutagenized treatment, selection treatment, and generation as fixed effects. Random effects were fit with block, replicate lineage nested with selection treatment, and starting males nested within block, replicate lineage, and selection treatment using glmmTMB (v0.2.3).

### Environmental Canalization

All artificially selected and genetically assimilated lineages were raised in low-density vials (n=3 vials each) at 6 different temperatures: 31°C, 29°C, 24.5°C, 21°C, 18°C, and 16°C. Adult flies were collected when they eclosed. Animals were stored in 70% EtOH and right wings were dissected from both males and females and imaged as described above (pleiotropy). Landmark and semi-landmark data was recorded in the same way, with the exception that Wings (v. 4.11.22) was used to fit splines. A total of 2870 individuals (females) were included, an average of 53.1 wings for each replicate lineage/temperature.

Wings of both sexes were scored for presence of posterior crossveins in addition to other wing perturbations (anterior cross vein loss, wing margin perturbation, longitudinal vein loss, additional veins). A total of 6346 individuals were scored (~average 58.7 wings sex/lineage/temperature). Proportions of posterior crossvein loss was calculated for each vial replicate within sex/lineage/temperature and modeled using a logistic mixed model with treatment, temperature, sex and their second-order interactions fit as fixed effects and a binomial distribution. A random effect of replicate lineage for each temperature:sex term was included in the model. Because of extremely low observed numbers for qualitative wing defects other than posterior cross vein loss, all other phenotypes were grouped into a single category (essentially wild type VS non wild type wing morphology). The proportion of wings with phenotypes were calculated and modeled as described above.

## Supporting Information

**S1 Figure:**
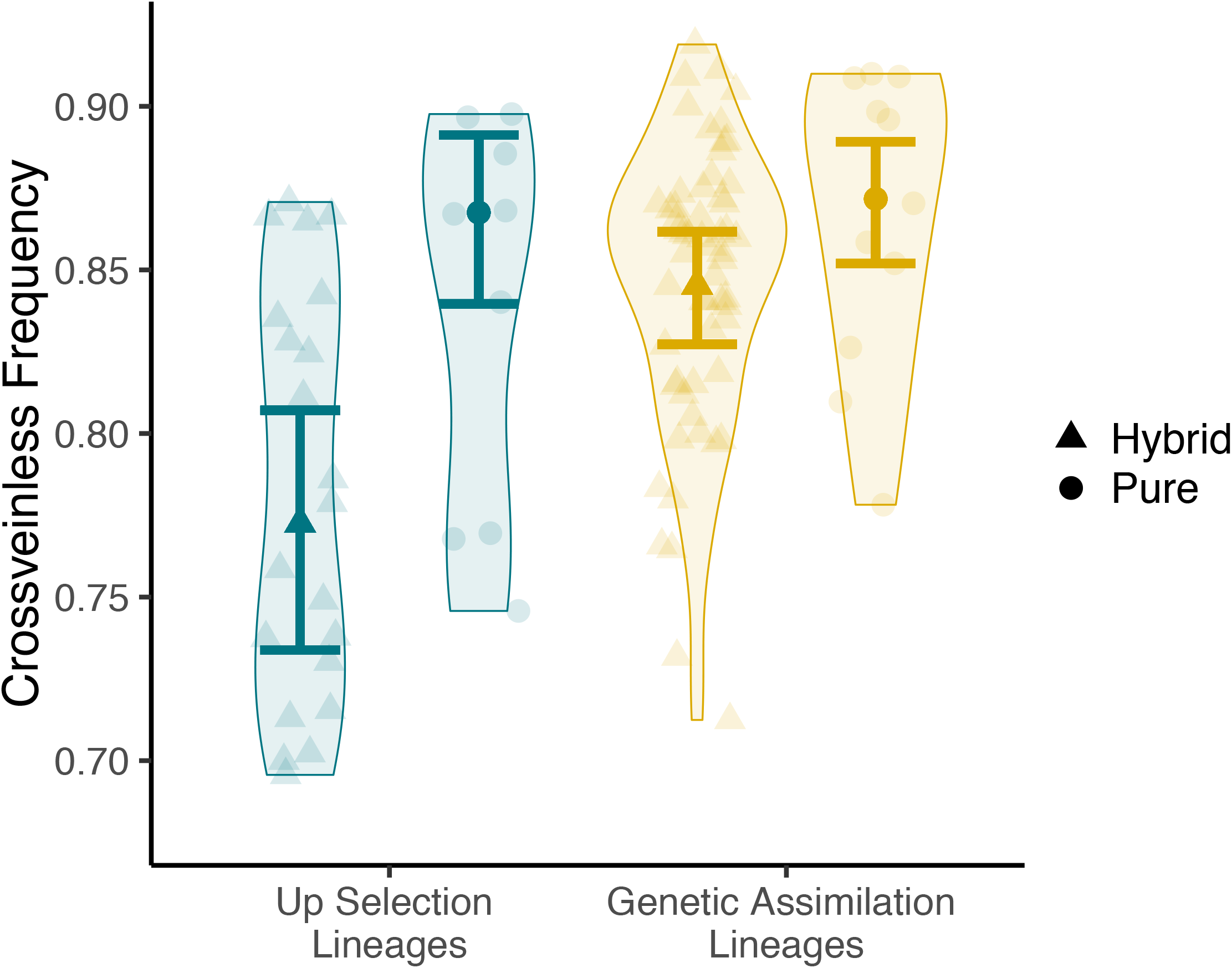
Incomplete parallelism among lineages in response to artificial selection or genetic assimilation. Partial di-allele crosses between the replicate genetically assimilated lineages were performed. Pure populations are those with parents from the same replicate lineage. Hybrid populations are those with parents from different replicate lineages. Crosses were done over 2 generational timepoints. Up selection lineages: nhybrid=24, npure=12. Genetic assimilation lineages: nhybrid=60, npure=12. Transparent points show variation and opaque points are fitted values for each set of crosses. Error bars are 95% significantly by the purity of the cross (p<0.05).

**S2 Figure:**
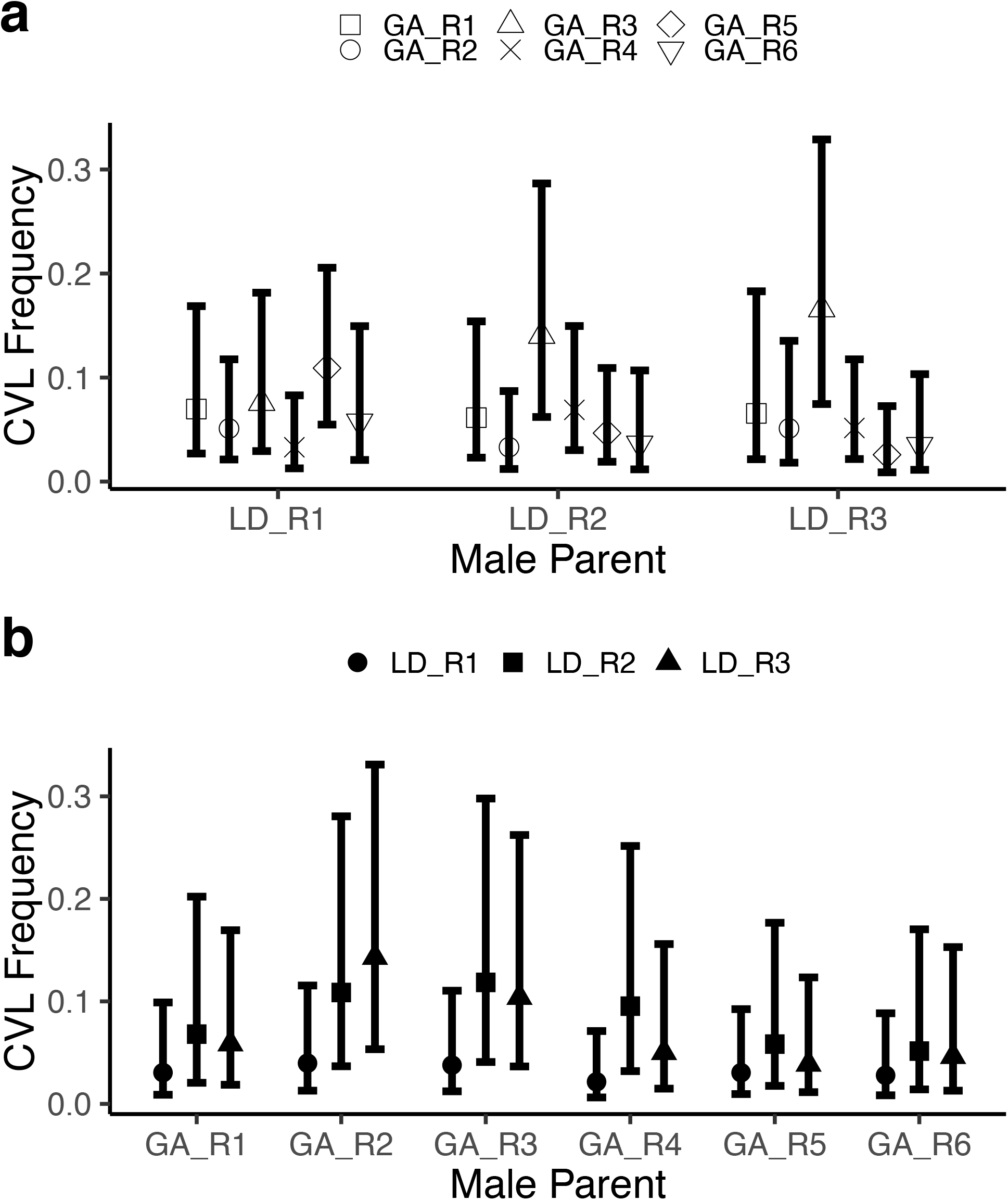
Low crossveinless frequency among hybrid progeny of genetic assimilation and lab domestication lineages. a) Male lab domestication and female genetic assimilation parents and b) the reciprocal male genetic assimilation and female lab domestication parent lineages (n=4). Error bars are 95% CIs on estimated effects from a generalized linear mixed model.

**S3 Figure:**
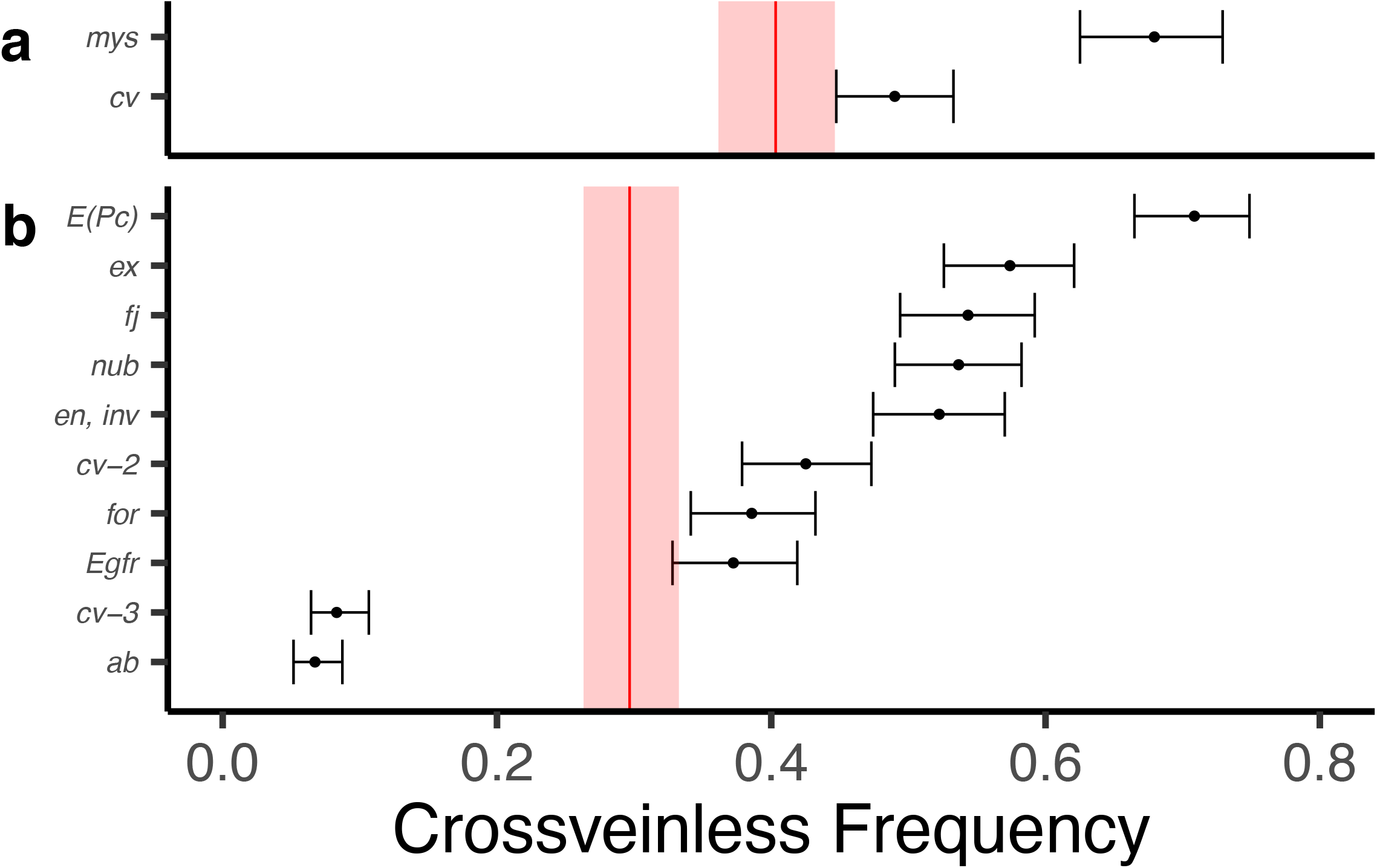
Deletion line crosses average for all genetic assimilation lineages in second experiment for further testing. Genes of interest in each deletion region for (a) Exelixis X chromosomes and (b) Exelixis autosomes (n=24 crosses for each deletion). Red solid lines represent Exelixis progenitor means with shaded rectangles as 95% CIs. Error bars are 95% CIs on estimated effects from a generalized linear mixed model.

**S4 Figure:**
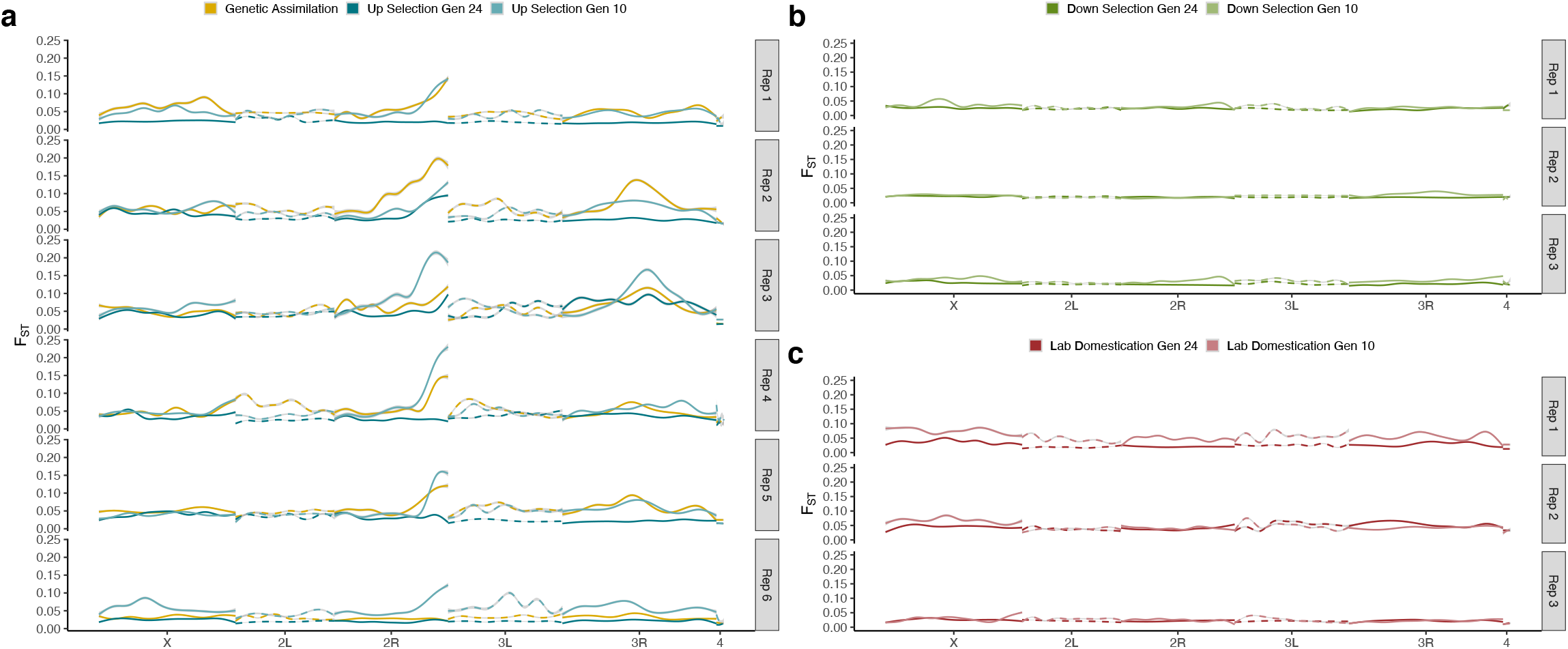
F_ST_ for (a) up-selection, (b) down-selection, and (c) lab domestication lineages at two time points during artificial selection and genetic assimilation lineages. F_ST_ was calculated using PoPoolation2 (Kofler et al. 2011b) on 500 base-pair windows in comparison with the ancestral population for two separate mapping software and taking the minimum F_ST_ value. Plots were generated with geom_smooth() using method ‘gam’ and formula ‘y~s(x,bs=“cs”)’ in R.

**S5 Figure:**
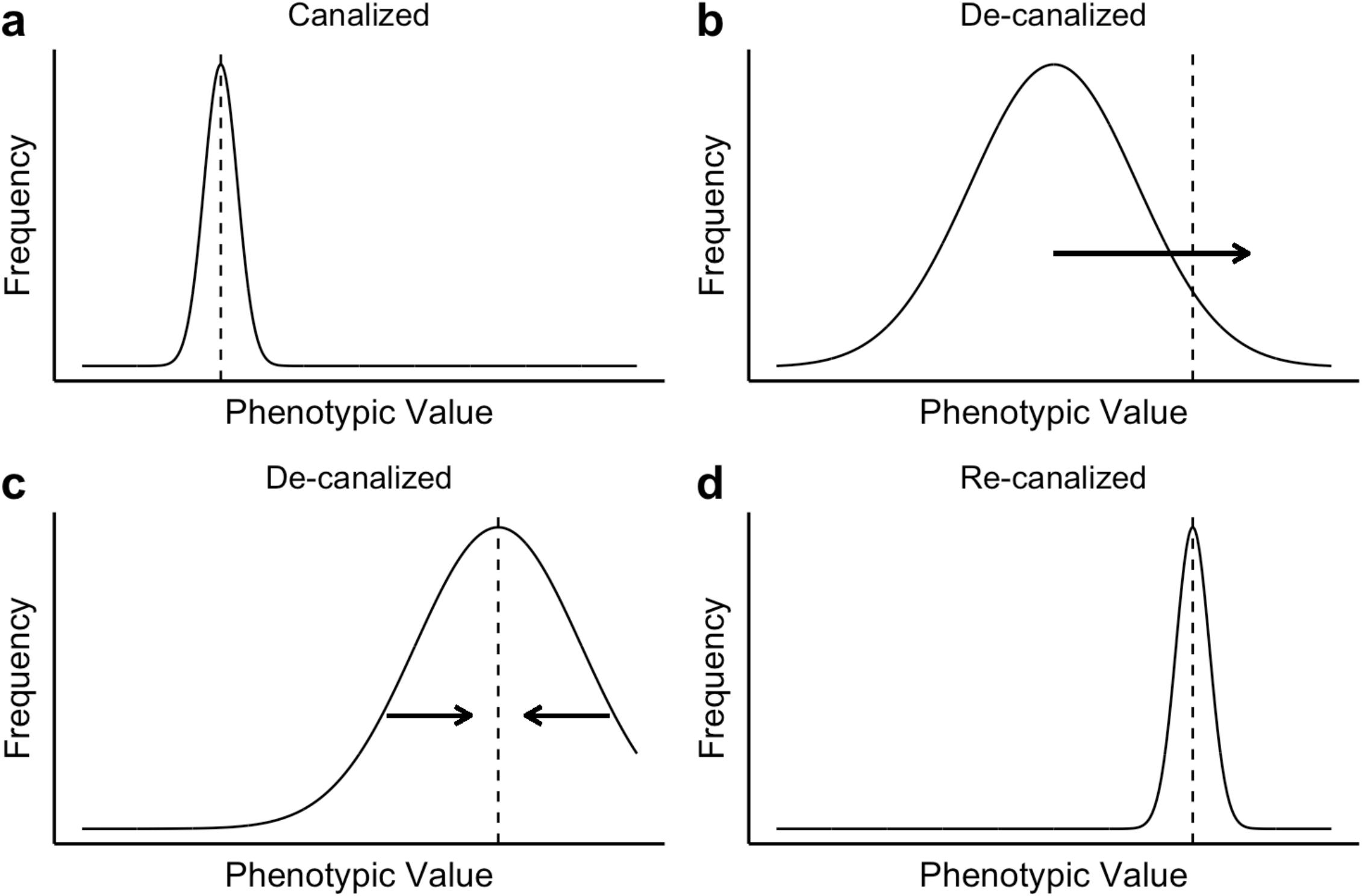
Waddington’s canalization model explaining genetic assimilation. (a) Canalaized system with phenotypic values distributed around the optimum (represented by a dashed line). (b) De-canalized system, where a new optimum is introduced and results in the disruption of canalization. Arrow represents selection shifting phenotypic mean. (c) The distribution of phenotype shifts towards the new optimum and stabilizing selection occurs. (d) Re-canalized system with phenotypic values distributed around the new optimum.

**S6 Figure:**
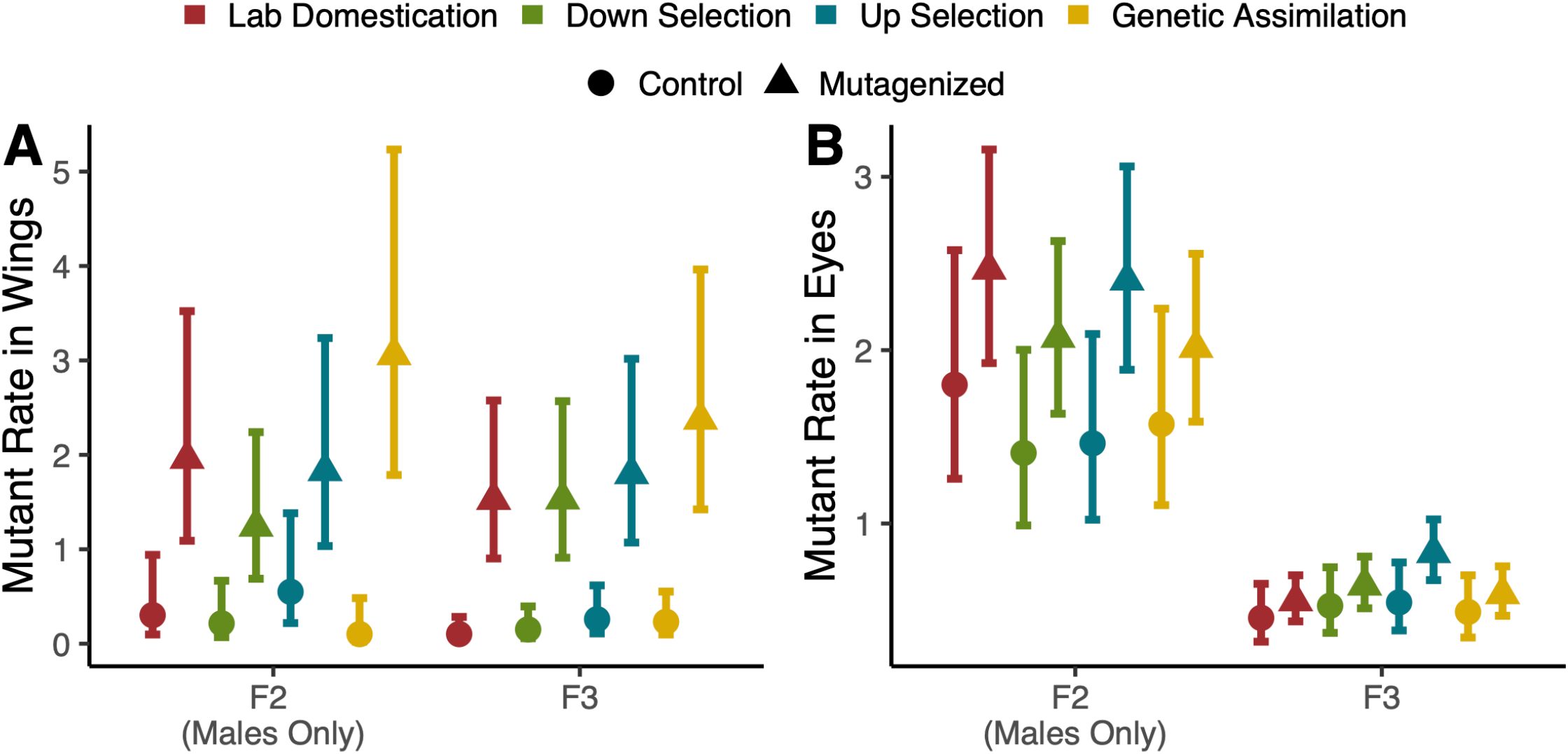
No differences in sensitivity to mutagenesis by way of mutant rate in (A) wings or (B) eyes (Mutagenesis Part 2). Mutant rate is the estimated rate of observed phenotypes per individual for either (A) wings or (B) eyes. n_Mutagenized_part2_ = 27-30 and n_Control_part2_ = 15 replicate vials sorted for each selection lineage. Error bars are 95% confidence intervals on estimated effects from a generalized linear mixed model.

**S7 Figure:**
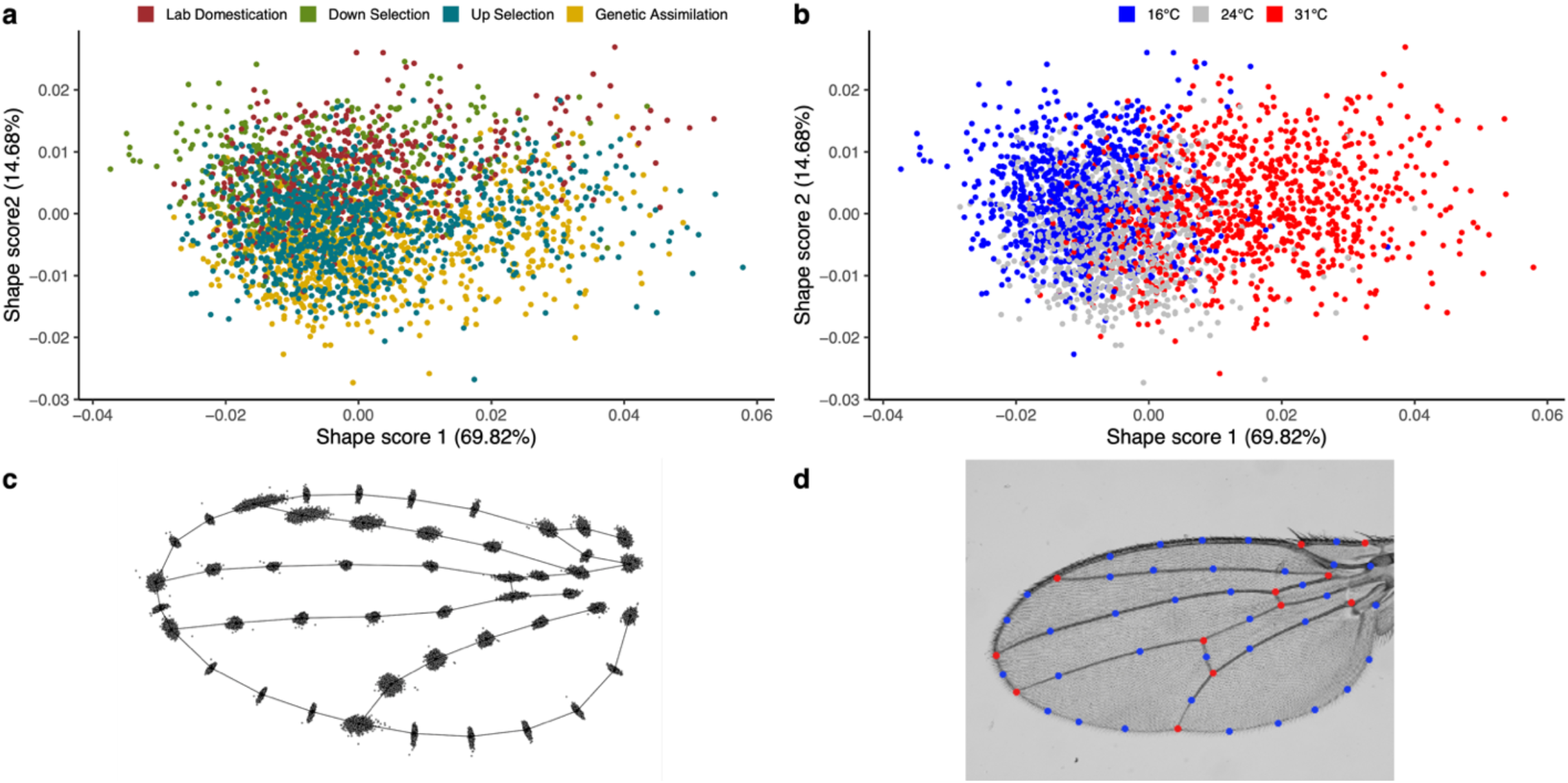
No differences in environmental canalization for wing shape among evolutionary lineages. **(a)** Shape scores for selection lineages and (b) same scores colored by rearing temperature show much of the variation in wing shape is due to temperature. Shape scores are projections of observed data onto vectors defined by PCs of fitted values. (c) All specimens’ landmark data plotted on a mean shape wing show small variation in distribution. (d) Depicts landmark positions on wing image.

**S8 Figure:**
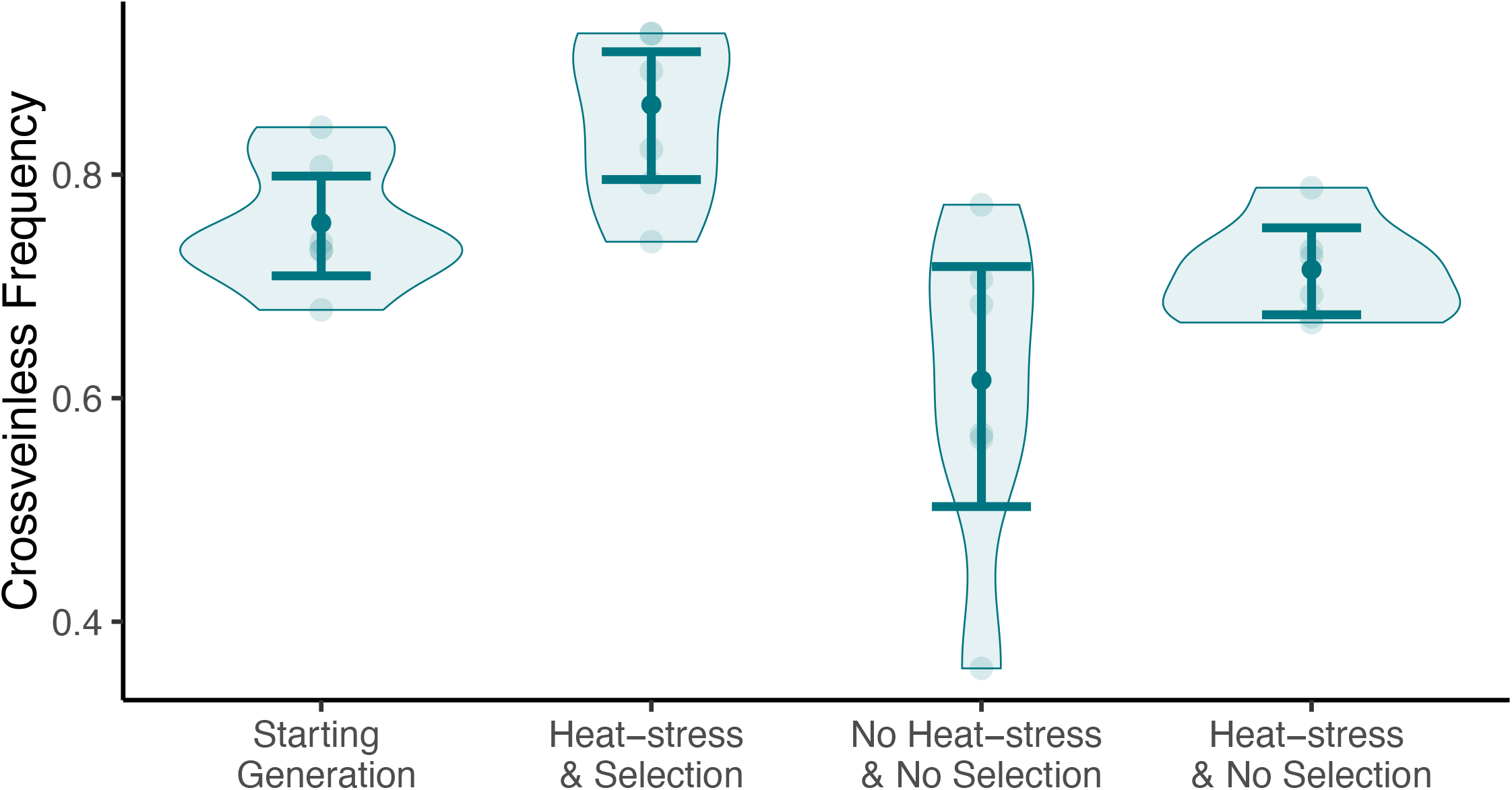
Relaxation of selection for five generations reduces the frequency of the CVL phenocopy response. Frequency of CVL after 5 generations of relaxed selection. The starting generation is the average CVL frequency of generations 17, 18, and 19. Heat-stress and selection are the lineages continued for the normal procedure of artificial selection. 200 individuals were counted for all six replicate lineages for each treatment. Transparent points show variation among replicate lineages and opaque points are fitted values for each treatment. Error bars are 95% confidence intervals. ANOVA from the generalized linear model shows that all treatments differed significantly from the starting generation, heat-stress & selection (p<0.0001), no heat-stress & no selection (p<0.0001) and heat-stress & no selection (p<0.01).

**S9 Figure:**
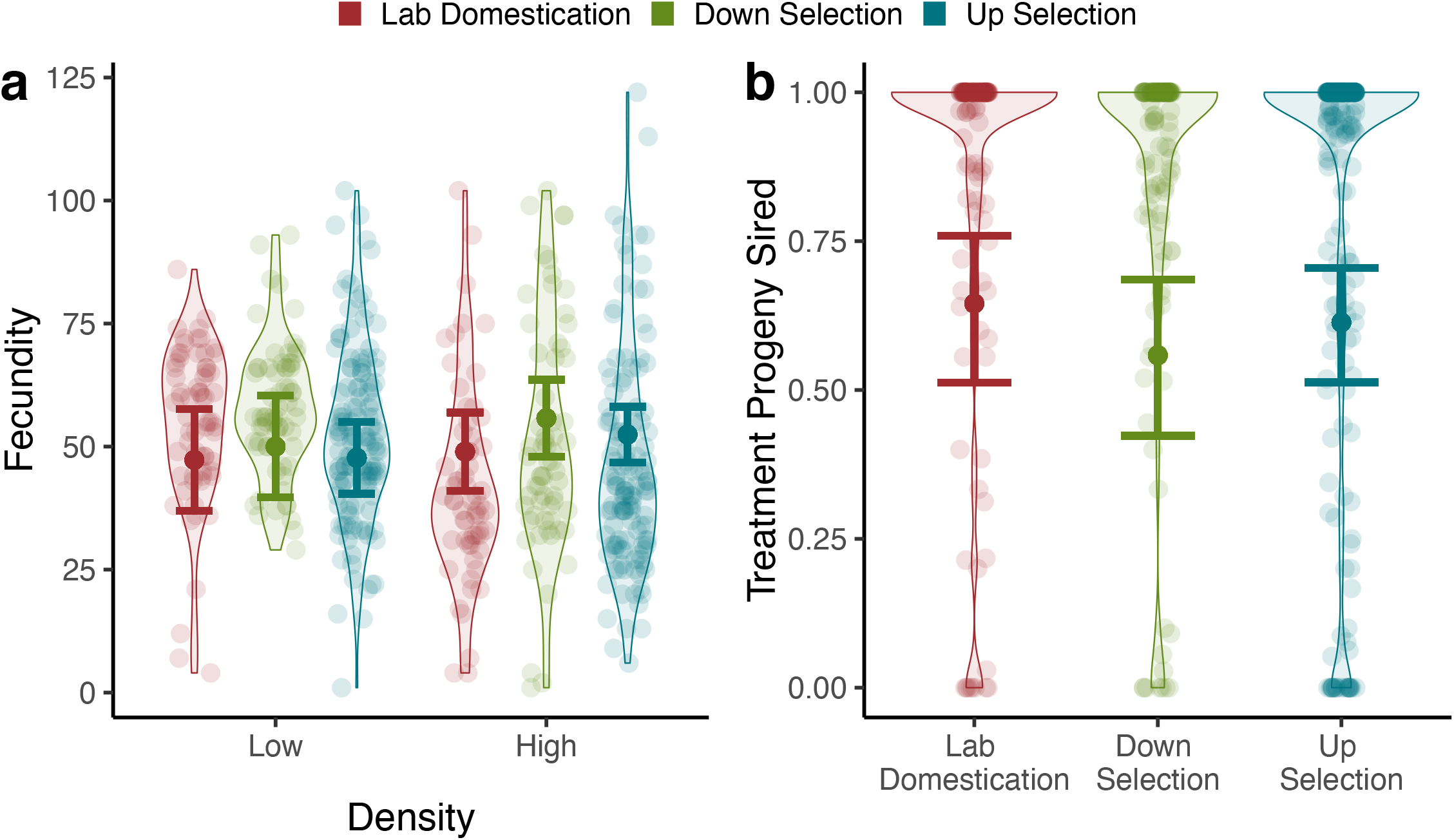
Variation in additional fitness components does not appear to be associated with frequency of CVL alleles. (a) Fecundity of the three selection regimes. Females were raised at either high or low densities before egg laying. For low density: n_LD_=67, n_DOWN_=68, n_UP_=138 females. For low density: n_LD_=62, n_DOWN_=64, n_UP_=126 females. (b) The observed proportion of treatment progeny for each selection lineage. n=37-50 per replicate lineage in a treatment. Competitor females were kept with a treatment and competitor male for 6 days and all progeny from that time was counted. Transparent points show variation within lineages and opaque points are fitted values for each lineage. Error bars are 95% confidence intervals on estimated effects from a generalized linear mixed model; none of the treatments show significant differences from each other at either density (p>0.1).

**S10 Figure:**
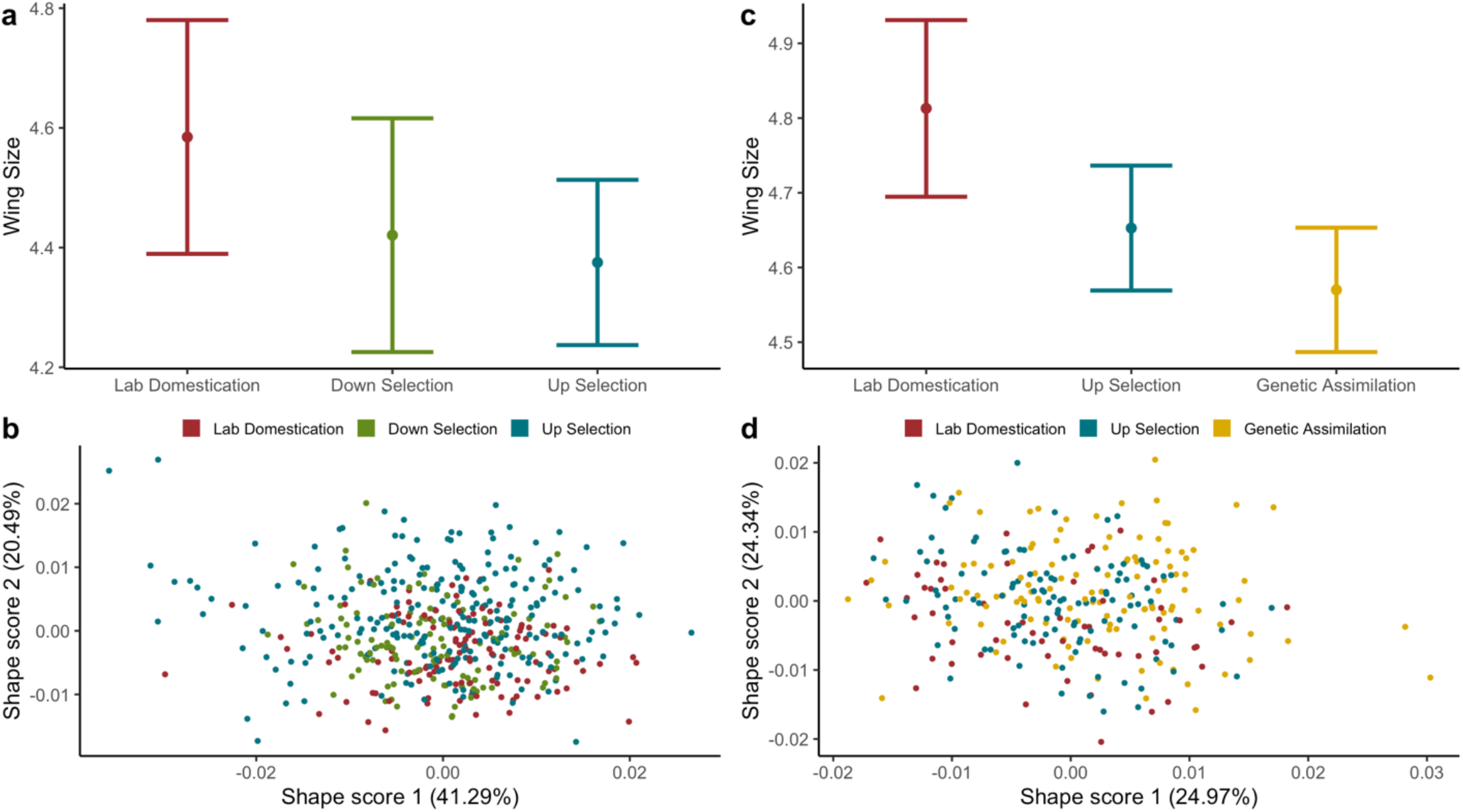
Variation in wing size and shape does not appear to be correlated with frequency of CVL alleles. Wing size (centroid size) and shape scores for two sets of comparisons among the selection lineages. The first group (a,b) was used for comparison between up-selection and down-selection (n~160/group), and the second group was (c,d) for up-selection and genetic assimilation (n~97/group). Error bars are 95% confidence intervals on estimated effects from a generalized linear mixed model. Levene’s statistic was used to estimate variability and showed no difference between treatment for either set of selection lineages. Shape scores are projections of observed data onto vectors defined by PCs of fitted values.

**S11 Figure:**
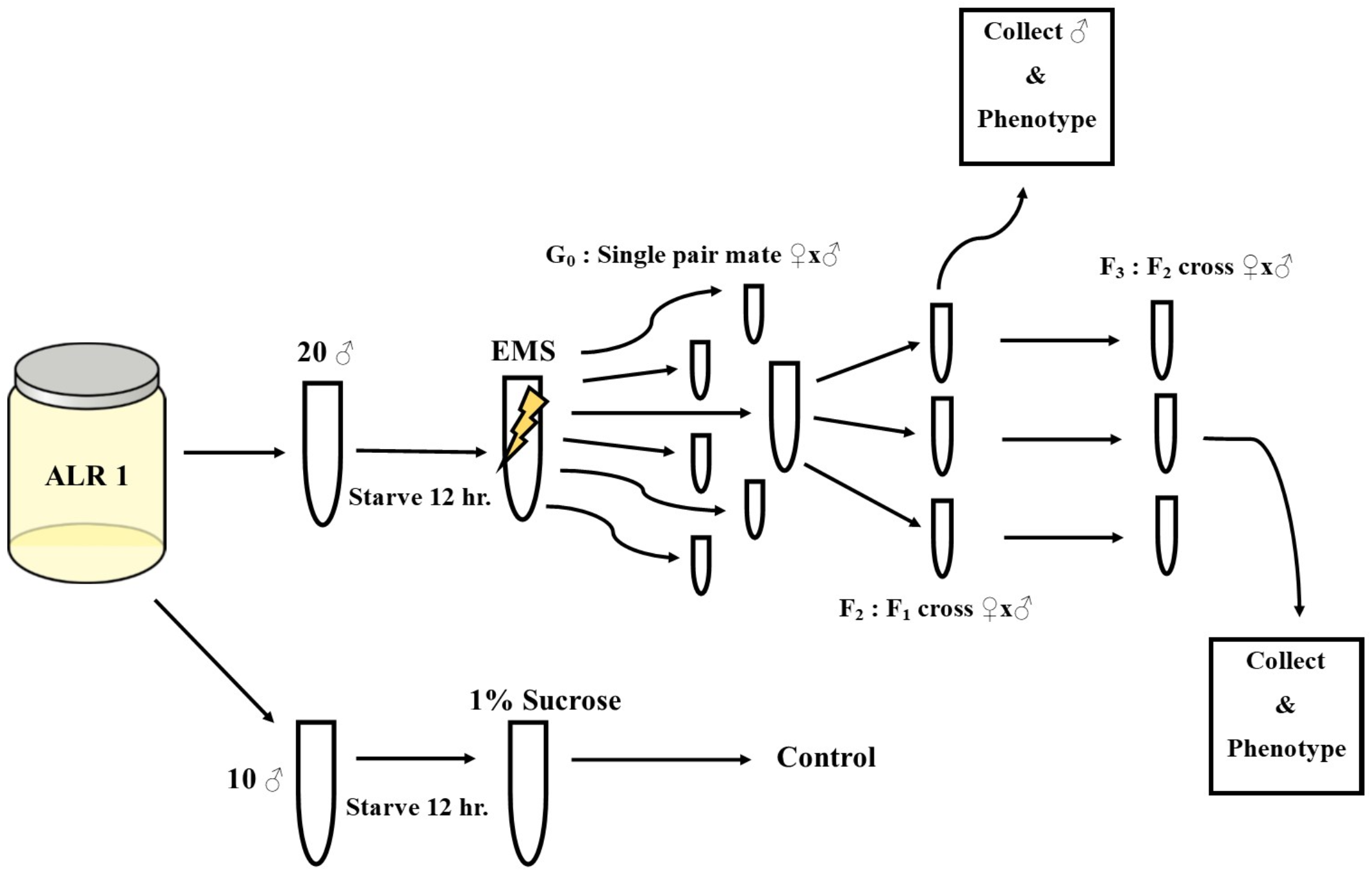
Graphic of methods for mutagenesis assay. For each genetic assimilated lineage (here “ALR 1” used as an example) 20 males were starved, exposed to EMS, and used in single mate pairing to create lines. Male progeny from generation 2 were scored for X-chromosome linked phenotypes and all progeny from generation 3 were scored for phenotypes. For each genetic assimilation lineage some males were used as a control, exposed to a sucrose solution, but otherwise scored the same.

**S1 Table:** Proportion of selected variants that are segregating in the ancestor. **All** variants that had an F_ST_ higher than 0.3 (F_ST_ used was minimum F_ST_ between two mappers, novoalign and BWA, calculated using PoPoolation2) were checked to see if present in the ancestral population. Major allele is allele with higher frequency than minor allele.

| Genetic Assimilation Replicate Lineage | Major allele is segregating | Major allele is new mutation | Minor allele is segregating | Minor allele is new mutation | Proportion of major allele segregating | Proportion of minor allele segregating |
| --- | --- | --- | --- | --- | --- | --- |
| 1 | 22590 | 495 | 19365 | 55 | 97.86% | 99.72% |
| 2 | 15669 | 146 | 11576 | 51 | 99.08% | 99.56% |
| 3 | 53986 | 1006 | 45231 | 80 | 98.17% | 99.82% |
| 4 | 26701 | 535 | 24757 | 163 | 98.04% | 99.35% |
| 5 | 21753 | 266 | 18191 | 69 | 98.79% | 99.62% |
| 6 | 17659 | 213 | 13011 | 93 | 98.81% | 99.29% |

**S2 Table:** Top 20+ largest regions for reduction in nucleotide diversity (Tajima’s pi). Reductions in Tajima’s pi were defined for each genetic assimilation lineages when showing a larger reduction from the ancestral pi as compared to the lab domestication lineages. **Top 20+ largest regions for reduction in nucleotide diversity (Nucleotide Diversity/Tajima’s π).**

| Genetic Assimilation Replicate 1 |  | Genetic Assimilation Replicate 2 |  | Genetic Assimilation Replicate 3 |  |
| --- | --- | --- | --- | --- | --- |
| Chromosome:Position | Size (Mb) | Chromosome:Position | Size (Mb) | Chromosome:Position | Size (Mb) |
| 2R:20171250-20188750 | 0.0175 | 2R:20370250-20386250 | 0.016 | 2R:18702750-18783250 | 0.0805 |
| 2R:20370250-20386250 | 0.016 | X:21377750-21393250 | 0.0155 | 2R:18658750-18691250 | 0.0325 |
| 2R:20027750-20043250 | 0.0155 | 2R:20027750-20043250 | 0.0155 | 3R:10808250-10840250 | 0.032 |
| 3L:22444750-22459750 | 0.015 | 2R:20124250-20137750 | 0.0135 | 3L:3163750-3189750 | 0.026 |
| 2R:20124250-20137750 | 0.0135 | 2R:20415750-20429250 | 0.0135 | 2R:20323750-20349750 | 0.026 |
| 2R:20415750-20429250 | 0.0135 | 2R:15545750-15558250 | 0.0125 | 2R:19470750-19494250 | 0.0235 |
| 2R:19565750-19578250 | 0.0125 | 2L:22339250-22351250 | 0.012 | 2R:20192250-20215750 | 0.0235 |
| 2R:20204250-20215750 | 0.0115 | 2R:20338250-20350250 | 0.012 | 3L:9608250-9630750 | 0.0225 |
| 3R:9241750-9252250 | 0.0105 | 3R:5540250-5551750 | 0.0115 | 2R:17312750-17334250 | 0.0215 |
| 3R:9714250-9724750 | 0.0105 | 3R:9241250-9252750 | 0.0115 | 3R:16445750-16467250 | 0.0215 |
| X:11705750-11714750 | 0.009 | 2R:20171250-20182250 | 0.011 | 2R:19372750-19393750 | 0.021 |
| X:14743250-14752250 | 0.009 | 2R:20387250-20397750 | 0.0105 | 2R:13762250-13782750 | 0.0205 |
| X:20912750-20921750 | 0.009 | 2L:19727250-19737250 | 0.01 | 2R:18273750-18294250 | 0.0205 |
| 3L:22708250-22717250 | 0.009 | 3L:4466250-4476250 | 0.01 | 2R:20083750-20104250 | 0.0205 |
| 3L:24044750-24053750 | 0.009 | 2R:18005250-18015250 | 0.01 | 2R:17948250-17968250 | 0.02 |
| 2R:20104750-20113750 | 0.009 | X:13004750-13013750 | 0.009 | 3R:10634250-10654250 | 0.02 |
| 2R:20151250-20160250 | 0.009 | X:14965250-14974250 | 0.009 | 2R:19660250-19679750 | 0.0195 |
| 2R:20387250-20396250 | 0.009 | 2R:1517750-1526750 | 0.009 | 2R:20494750-20513750 | 0.019 |
| 3R:9645250-9654250 | 0.009 | 2R:17347250-17356250 | 0.009 | 2R:18421750-18440250 | 0.0185 |
| 3R:9756750-9765750 | 0.009 | X:7620250-7628750 | 0.0085 | 2R:19506250-19524750 | 0.0185 |
|  |  | X:17453250-17461750 | 0.0085 |  |  |
|  |  | 2R:16191750-16200250 | 0.0085 |  |  |
|  |  | 2R:16712750-16721250 | 0.0085 |  |  |
|  |  | 2R:20147250-20155750 | 0.0085 |  |  |
|  |  | 2R:20482250-20490750 | 0.0085 |  |  |
|  |  | 3R:20845250-20853750 | 0.0085 |  |  |

**S2 Table (Cont.): Top 20+ largest regions for reduction in nucleotide diversity (Nucleotide Diversity/Tajima's $\pi$ ).**
| Genetic Assimilation Replicate 4 |  | Genetic Assimilation Replicate 5 |  | Genetic Assimilation Replicate 6 |  |
| --- | --- | --- | --- | --- | --- |
| Chromosome:Position | Size (Mb) | Chromosome:Position | Size (Mb) | Chromosome:Position | Size (Mb) |
| 2R:14515250-14529750 | 0.0145 | 2R:20120250-20217750 | 0.0975 | 2R:20171250-20198250 | 0.027 |
| 3R:11438250-11452250 | 0.014 | 2R:20370250-20413250 | 0.043 | 3L:22497250-22518750 | 0.0215 |
| 2R:17343250-17356250 | 0.013 | 2R:20288250-20322250 | 0.034 | 2R:20338250-20354750 | 0.0165 |
| 2R:12887250-12898250 | 0.011 | 2R:20331750-20354750 | 0.023 | 2R:20370250-20386250 | 0.016 |
| 3R:14682750-14693750 | 0.011 | 2R:20098250-20119250 | 0.021 | 2R:20123250-20137750 | 0.0145 |
| 3R:10674750-10685250 | 0.0105 | 2R:20417750-20431750 | 0.014 | 2R:19285750-19299750 | 0.014 |
| 3R:15424250-15434750 | 0.0105 | 2R:20355750-20369250 | 0.0135 | 2R:20146250-20160250 | 0.014 |
| 3L:10295750-10305750 | 0.01 | 2R:20218750-20231750 | 0.013 | 3L:22446250-22458750 | 0.0125 |
| 2R:17167250-17177250 | 0.01 | 2R:20274250-20287250 | 0.013 | 2R:19223750-19235750 | 0.012 |
| 3R:12176750-12186750 | 0.01 | 2R:20251250-20263250 | 0.012 | 2R:20398750-20410750 | 0.012 |
| 3R:17418250-17428250 | 0.01 | X:22036250-22047250 | 0.011 | 2R:20204250-20215750 | 0.0115 |
| 2R:18005750-18015250 | 0.0095 | 2R:19405750-19416750 | 0.011 | 3R:9241750-9252750 | 0.011 |
| 2L:18019750-18028750 | 0.009 | 3L:3169750-3180250 | 0.0105 | 2R:18004250-18014750 | 0.0105 |
| 2L:20200750-20209750 | 0.009 | X:20912250-20922250 | 0.01 | 2R:20387250-20397750 | 0.0105 |
| 3R:15482750-15491750 | 0.009 | 2R:19223750-19233750 | 0.01 | 2L:19750750-19760750 | 0.01 |
| 3R:2940250-2948750 | 0.0085 | 2R:20470250-20480250 | 0.01 | 3L:24044750-24054750 | 0.01 |
| 3R:7705250-7713750 | 0.0085 | 3R:13062750-13072250 | 0.0095 | 2L:22342750-22352250 | 0.0095 |
| 3R:10550250-10558750 | 0.0085 | X:21388250-21397250 | 0.009 | 3L:24062250-24071750 | 0.0095 |
| 3R:10570750-10579250 | 0.0085 | 2R:17051750-17060750 | 0.009 | 2R:17345250-17354750 | 0.0095 |
| 2L:18609750-18617750 | 0.008 | 2R:18006750-18015750 | 0.009 | 2R:20422250-20431750 | 0.0095 |
| 2R:16705250-16713250 | 0.008 | 3R:10634750-10643750 | 0.009 |  |  |
| 3R:7646250-7654250 | 0.008 |  |  |  |  |
| 3R:10734750-10742750 | 0.008 |  |  |  |  |
| 3R:12051750-12059750 | 0.008 |  |  |  |  |
| 3R:15334250-15342250 | 0.008 |  |  |  |  |

**S3 Table:**
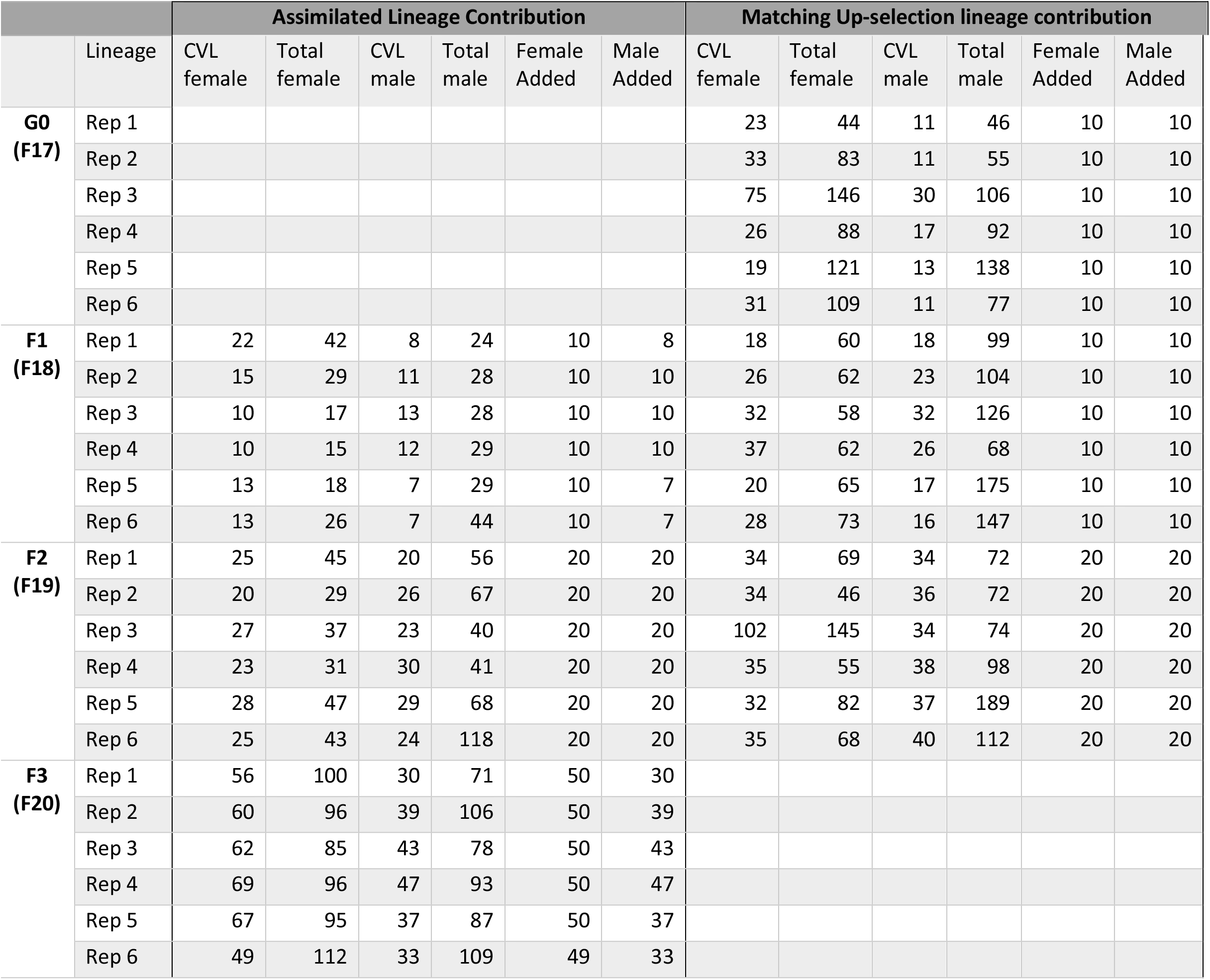
Supplemented assimilated flies to each genetically assimilated lineage from the corresponding up-selection lineage. Each genetically assimilated lineages was started (G0) from its corresponding up-selection lineage in generation 17 of the artificial selection experiment. For generations 1 and 2 (corresponding to F18 and F19 of up-selection), assimilated males and females were combined from the genetic assimilated lineage and corresponding up-selection lineage to continue the next generation. By generation 3, we had enough assimilated individuals to sustain the genetically assimilated lineages independently.

|  |  | Assimilated Lineage Contribution |  |  |  |  |  | Matching Up-selection lineage contribution |  |  |  |  |  |
| --- | --- | --- | --- | --- | --- | --- | --- | --- | --- | --- | --- | --- | --- |
|  | Lineage | CVL female | Total female | CVL male | Total male | Female Added | Male Added | CVL female | Total female | CVL male | Total male | Female Added | Male Added |
| **G0 (F17)** | Rep 1 |  |  |  |  |  |  | 23 | 44 | 11 | 46 | 10 | 10 |
|  | Rep 2 |  |  |  |  |  |  | 33 | 83 | 11 | 55 | 10 | 10 |
|  | Rep 3 |  |  |  |  |  |  | 75 | 146 | 30 | 106 | 10 | 10 |
|  | Rep 4 |  |  |  |  |  |  | 26 | 88 | 17 | 92 | 10 | 10 |
|  | Rep 5 |  |  |  |  |  |  | 19 | 121 | 13 | 138 | 10 | 10 |
|  | Rep 6 |  |  |  |  |  |  | 31 | 109 | 11 | 77 | 10 | 10 |
| **F1 (F18)** | Rep 1 | 22 | 42 | 8 | 24 | 10 | 8 | 18 | 60 | 18 | 99 | 10 | 10 |
|  | Rep 2 | 15 | 29 | 11 | 28 | 10 | 10 | 26 | 62 | 23 | 104 | 10 | 10 |
|  | Rep 3 | 10 | 17 | 13 | 28 | 10 | 10 | 32 | 58 | 32 | 126 | 10 | 10 |
|  | Rep 4 | 10 | 15 | 12 | 29 | 10 | 10 | 37 | 62 | 26 | 68 | 10 | 10 |
|  | Rep 5 | 13 | 18 | 7 | 29 | 10 | 7 | 20 | 65 | 17 | 175 | 10 | 10 |
|  | Rep 6 | 13 | 26 | 7 | 44 | 10 | 7 | 28 | 73 | 16 | 147 | 10 | 10 |
| **F2 (F19)** | Rep 1 | 25 | 45 | 20 | 56 | 20 | 20 | 34 | 69 | 34 | 72 | 20 | 20 |
|  | Rep 2 | 20 | 29 | 26 | 67 | 20 | 20 | 34 | 46 | 36 | 72 | 20 | 20 |
|  | Rep 3 | 27 | 37 | 23 | 40 | 20 | 20 | 102 | 145 | 34 | 74 | 20 | 20 |
|  | Rep 4 | 23 | 31 | 30 | 41 | 20 | 20 | 35 | 55 | 38 | 98 | 20 | 20 |
|  | Rep 5 | 28 | 47 | 29 | 68 | 20 | 20 | 32 | 82 | 37 | 189 | 20 | 20 |
|  | Rep 6 | 25 | 43 | 24 | 118 | 20 | 20 | 35 | 68 | 40 | 112 | 20 | 20 |
| **F3 (F20)** | Rep 1 | 56 | 100 | 30 | 71 | 50 | 30 |  |  |  |  |  |  |
|  | Rep 2 | 60 | 96 | 39 | 106 | 50 | 39 |  |  |  |  |  |  |
|  | Rep 3 | 62 | 85 | 43 | 78 | 50 | 43 |  |  |  |  |  |  |
|  | Rep 4 | 69 | 96 | 47 | 93 | 50 | 47 |  |  |  |  |  |  |
|  | Rep 5 | 67 | 95 | 37 | 87 | 50 | 37 |  |  |  |  |  |  |
|  | Rep 6 | 49 | 112 | 33 | 109 | 49 | 33 |  |  |  |  |  |  |

## References

1. Waddington CH. Canalization of development and the inheritance of acquired characters. Nature. 1942;150: 563–565. doi:10.1038/150405d0

2. Waddington CH. Genetic assimilation of an acquired character. Evolution. 1953;7: 118–126.

3. Mohler JD. Preliminary genetic analysis of crossveinless-like strains of Drosophila melanogaster. Genetics. 1965;51: 641–651.

4. Milkman RD. The genetic basis of natural variation. V. Selection for crossveinless polygenes in new wild strains of Drosophila melanogaster. Genetics. 1964;50: 625–632.

5. Milkman RD. The genetic basis of natural variation. I. Crossveins in Drosophila melanogaster. Genetics. 1960;45: 35–48.

6. Bateman KG. The genetic assimilation of four venation phenocopies. Journal of genetics. 1959;56: 443–474.

7. Milkman RD. The genetic basis of natural variation. II. Analysis of a polygenic system in Drosophila melanogaster. Genetics. 1960;45: 377–391.

8. Paaby AB, White AG, Riccardi DD, Gunsalus KC, Piano F, Rockman M v. Wild worm embryogenesis harbors ubiquitous polygenic modifier variation. eLife. 2015;4: e09178. doi:10.7554/eLife.09178

9. Ledón-Rettig Cc, Pfennig DW, Crespi EJ. Diet and hormonal manipulation reveal cryptic genetic variation: implications for the evolution of novel feeding strategies. Proceedings Biological sciences / The Royal Society. 2010;277: 3569–78. doi:10.1098/rspb.2010.0877

10. Aubret F, Shine R. Genetic Assimilation and the Postcolonization Erosion of Phenotypic Plasticity in Island Tiger Snakes. Current Biology. 2009;19: 1932–1936. doi:10.1016/j.cub.2009.09.061

11. Heil M, Greiner S, Meimberg H, Krüger R, Noyer J-L, Heubl G, et al. Evolutionary change from induced to constitutive expression of an indirect plant resistance. Nature. 2004;430: 205–208. doi:10.1038/nature02703

12. Ryan CP, Brownlie JC, Whyard S. Hsp90 and physiological stress are linked to autonomous transposon mobility and heritable genetic change in nematodes. Genome Biology and Evolution. 2016;8: 3794–3805. doi:10.1093/gbe/evw284

13. Swaegers J, Spanier KI, Stoks R. Genetic compensation rather than genetic assimilation drives the evolution of plasticity in response to mild warming across latitudes in a damselfly. Molecular Ecology. 2020;29: 4823–4834. doi:10.1111/mec.15676

14. Lack JB, Monette MJ, Johanning EJ, Sprengelmeyer QD, Pool JE. Decanalization of wing development accompanied the evolution of large wings in high-altitude Drosophila. Proceedings of the National Academy of Sciences. 2016;113: 1014–1019. doi:10.1073/pnas.1515964113

15. Landauer W. On phenocopies, their developmental physiology and genetic meaning. The American Naturalist. 1958;92: 201–213.

16. Fanti L, Piacentini L, Cappucci U, Casale AM, Pimpinelli S. Canalization by selection of de novo induced mutations. Genetics. 2017;206: 1995–2006. doi:10.1534/genetics.117.201079

17. van der Burg Krl, Lewis JJ, Brack BJ, Fandino RA, Mazo-Vargas A, Reed RD. Genomic architecture of a genetically assimilated seasonal color pattern. Science. 2020;370: 721–725. doi:10.1126/science.aaz3017

18. Vigne P, Gimond C, Ferrari C, Vielle A, Hallin J, Pino-Querido A, et al. A single-nucleotide change underlies the genetic assimilation of a plastic trait. Science Advances. 2021;7: 1–13. doi:10.1126/sciadv.abd9941

19. Duveau F, Félix M-A. Role of Pleiotropy in the Evolution of a Cryptic Developmental Variation in Caenorhabditis elegans. Noor Maf, editor. PLoS Biology. 2012;10: e1001230. doi:10.1371/journal.pbio.1001230

20. Milkman RD. The genetic basis of natural variation. IV. On the natural distribution of cve Polygenes of Drosophila melanogaster. Genetics. 1962;47: 261–272.

21. Milkman RD. The genetic basis of natural variation. III. Developmental lability and evolutionary potential. Genetics. 1961;46: 25–38.

22. Rendel JM. Canalization of the scute phenotype of Drosophila. Evolution. 1959;13: 425–439.

23. Waddington CH. Genetic Assimilation. Advances in Genetics. Academic Press; 1961. pp. 257–293. doi:10.1016/S0065-2660(08)60119-4

24. Boyer B, Parris D, Milkman RD. The Crossveinless Polygenes in an Iowa Population. Genetics. 1973;75: 169–179.

25. Ito H, Gaubert H, Bucher E, Mirouze M, Vaillant I, Paszkowski J. An siRNA pathway prevents transgenerational retrotransposition in plants subjected to stress. Nature. 2011;472: 115–119. doi:10.1038/nature09861

26. Cappucci U, Noro F, Casale AM, Fanti L, Berloco M, Alagia AA, et al. The Hsp70 chaperone is a major player in stress-induced transposable element activation. Proceedings of the National Academy of Sciences. 2019;116: 17943–17950. doi:10.1073/pnas.1903936116

27. Quadrana L, Etcheverry M, Gilly A, Caillieux E, Madoui M-A, Guy J, et al. Transposition favors the generation of large effect mutations that may facilitate rapid adaption. Nature Communications. 2019;10: 3421. doi:10.1038/s41467-019-11385-5

28. Specchia V, Piacentini L, Tritto P, Fanti L, Dalessandro R, Palumbo G, et al. Hsp90 prevents phenotypic variation by suppressing the mutagenic activity of transposons. Nature. 2010;463: 662–665. doi:10.1038/nature08739

29. Ryan CP, Brownlie JC, Whyard S. Hsp90 and physiological stress are linked to autonomous transposon mobility and heritable genetic change in nematodes. Genome Biology and Evolution. 2016;8: 3794–3805. doi:10.1093/gbe/evw284

30. Groth BR, Huang Y, Monette MJ, Pool JE. Directional selection reduces developmental canalization against genetic and environmental perturbations in drosophila wings. Evolution. 2018;72: 1708–1715. doi:10.1111/evo.13550

31. Masel J, Siegal ML. Robustness: mechanisms and consequences. Trends in Genetics. 2009;25: 395–403. doi:10.1016/j.tig.2009.07.005

32. Hermisson J, Wagner GP. The population genetic theory of hidden variation and genetic robustness. Genetics. 2004;168: 2271–2284. doi:10.1534/genetics.104.029173

33. Geiler-Samerotte K, Sartori FMO, Siegal ML. Decanalizing thinking on genetic canalization. Seminars in Cell and Developmental Biology. 2019;88: 54–66. doi:10.1016/j.semcdb.2018.05.008

34. Schrader L, Schmitz J. The impact of transposable elements in adaptive evolution. Molecular Ecology. 2019;28: 1537–1549. doi:10.1111/mec.14794

35. Dworkin I, Gibson G. Epidermal growth factor receptor and transforming growth factor-Beta signaling contributes to variation for wing shape in Drosophila melanogaster. Genetics. 2006;173: 1417–1431. doi:10.1534/genetics.105.053868

36. Debat V, Milton CC, Rutherford S, Klingenberg CP, Hoffmann A a. Hsp90 and the quantitative variation of wing shape in Drosophila melanogaster. Evolution; international journal of organic evolution. 2006;60: 2529–2538. doi:10.1111/j.0014-3820.2006.tb01887.x

37. Matsuda S, Shimmi O. Directional transport and active retention of Dpp / BMP create wing vein patterns in Drosophila. Developmental Biology. 2012;366: 153–162. doi:10.1016/j.ydbio.2012.04.009

38. Rohner N, Jarosz DF, Kowalko JE, Yoshizawa M, Jeffery WR, Borowsky RL, et al. Cryptic Variation in Morphological Evolution: HSP90 as a Capacitor for Loss of Eyes in Cavefish. Science. 2013;342: 1372–1375. doi:10.1126/science.1240276

39. Rutherford SL, Lindquist S. Hsp90 as a capacitor for morphological evolution. Nature. 1998;396: 336–342. doi:10.1038/24550

40. Deming D. Do extraordinary claims require extraordinary evidence? Philosophia. 2016;44: 1319–1331. doi:10.1007/s11406-016-9779-7

41. Kaufman M. First Contact: Scientific Breakthroughs in the Hunt for Life Beyond Earth. New York: Simon & Schuster; 2012.

42. Bainbridge SP, Bownes M. Staging the metamorphosis of Drosophila melanogaster. Journal of embryology and experimental morphology. 1981;66: 57–80. Available: http://www.ncbi.nlm.nih.gov/pubmed/6802923

43. Márquez EJ. CPR: using Drosophila wing shape data. 2012.

44. Adams DC, Collyer ML, Kaliontzopoulou A. Geomorph: software for geometric morphometric analyses. R package version 3.1.1. R Foundation for Statistical Computing. Vienna, Austria; 2019.

45. Kofler R, Langmüller AM, Nouhaud P, Otte KA, Schlötterer C. Suitability of different mapping algorithms for genome-wide polymorphism scans with pool-seq data. G3: Genes, Genomes, Genetics. 2016;6: 3507–3515. doi:10.1534/g3.116.034488

46. Kofler R, Orozco-terWengel P, de Maio N, Pandey RV, Nolte V, Futschik A, et al. Popoolation: A toolbox for population genetic analysis of next generation sequencing data from pooled individuals. PLoS ONE. 2011;6. doi:10.1371/journal.pone.0015925

47. Kofler R, Pandey RV, Schlötterer C. PoPoolation2: Identifying differentiation between populations using sequencing of pooled DNA samples (Pool-Seq). Bioinformatics. 2011;27: 3435–3436. doi:10.1093/bioinformatics/btr589

